# Catabolism of germinant amino acids is required to prevent premature spore germination in *Bacillus subtilis*

**DOI:** 10.1101/2024.02.22.581590

**Authors:** Iqra R. Kasu, Octavio Reyes-Matte, Alejandro Bonive-Boscan, Alan I. Derman, Javier Lopez-Garrido

## Abstract

Spores of *Bacillus subtilis* germinate in response to specific germinant molecules that are recognized by receptors in the spore envelope. Germinants signal to the dormant spore that the environment can support vegetative growth, so many germinants, such as alanine and valine, are also essential metabolites. As such, they are also required to build the spore. Here we show that these germinants cause premature germination if they are still present at the latter stages of spore formation and beyond, but that *B. subtilis* metabolism is configured to prevent this: alanine and valine are catabolized and cleared from wild-type cultures even when alternative carbon and nitrogen sources are present. Alanine and valine accumulate in the spent media of mutants that are unable to catabolize these amino acids, and premature germination is pervasive. Premature germination does not occur if the germinant receptor that responds to alanine and valine is eliminated, or if wild-type strains that are able to catabolize and clear alanine and valine are also present in coculture. Our findings demonstrate that spore-forming bacteria must fine-tune the concentration of any metabolite that can also function as a germinant to a level that is high enough to allow for spore development to proceed, but not so high as to promote premature germination. These results indicate that germinant selection and metabolism are tightly linked, and suggest that germinant receptors evolve in tandem with the catabolic priorities of the spore-forming bacterium.

## INTRODUCTION

Organisms can survive adverse conditions by reducing their metabolic activities (1–3). Bacteria such as *Bacillus subtilis* and other Firmicutes go so far as to produce metabolically dormant endospores (henceforth, spores) that are resistant to extreme environmental challenges including heat, starvation, desiccation, and antibiotics (4, 5). Spores are produced through a developmental program that entails the cooperation between two cells of unequal size and developmental fate (Fig. 1A): the smaller forespore, which will become the spore, and the larger mother cell, which lyses once spore formation is completed (6, 7). The forespore and mother cell originate from a cell division event in which the septum is formed close to one cell pole (8). The forespore is subsequently internalized within the mother cell cytoplasm (9), and matures in a process that involves the synthesis of several different layers that encase the spore (10–12), and the partial dehydration of the forespore core. Some of the water in the spore core is replaced by a chelate of calcium with a spore-specific molecule called dipicolinic acid (DPA). Forespore dehydration is essential for heat-resistance (5), and confers upon the spores their characteristic bright aspect under phase-contrast microscopy (Fig. 1B). After spore maturation, the mother cell lyses and the mature spore is released to the environment, where it can remain metabolically dormant for years (13).

**Figure 1.**
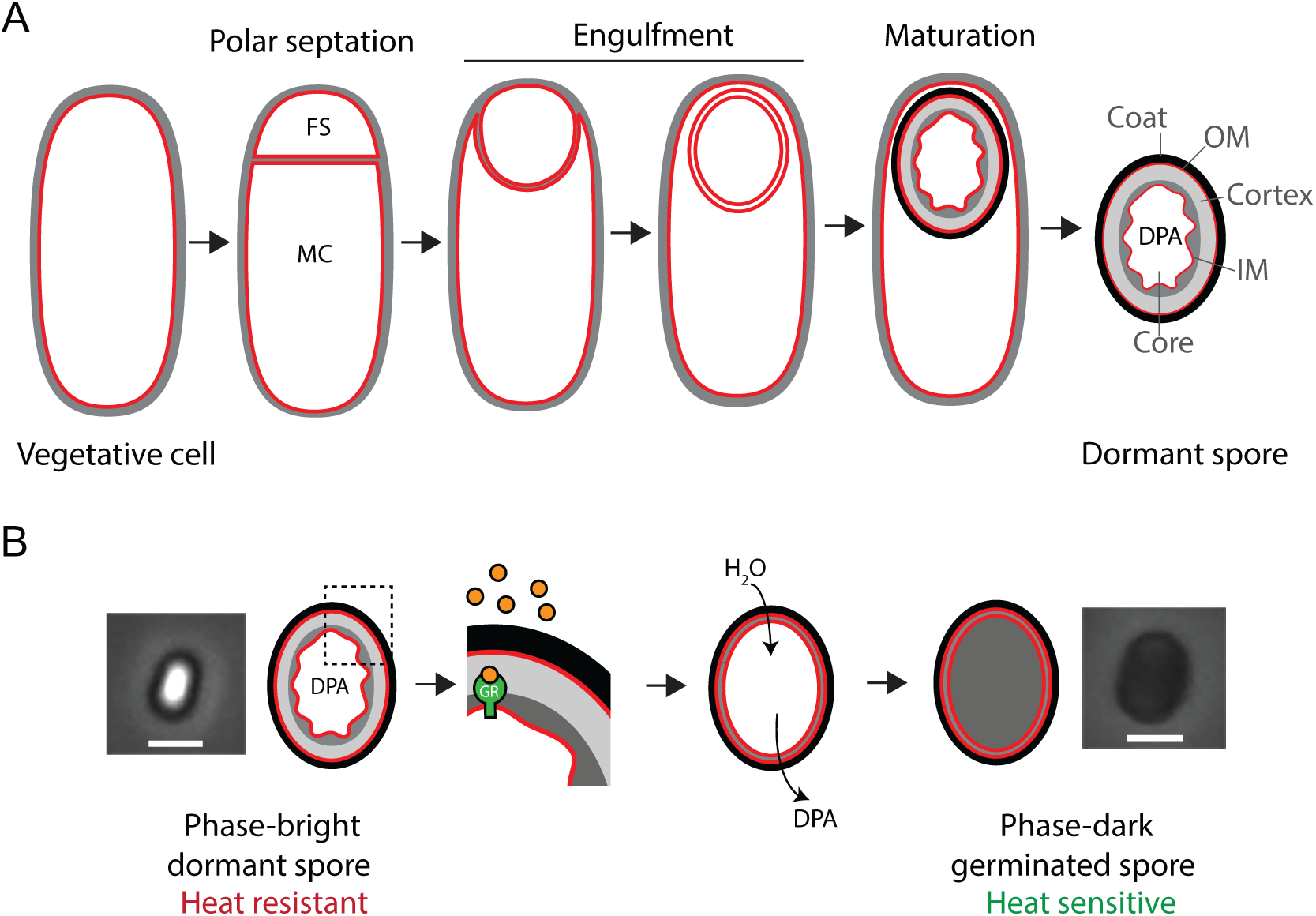
Sporulation and germination in *B. subtilis*. **(A)** Schematic of sporulation, with membranes in red and peptidoglycan in grey. Upon polar septation, the vegetative cell divides into a small forespore (FS) and a large mother cell (MC). The forespore is engulfed by the mother cell and matures within the mother cell cytoplasm. Maturation involves the formation of a peptidoglycan cortex (grey) and a proteinaceous coat (black) around the forespore, and the dehydration of the forespore core, which accumulates dipicolinic acid (DPA). Once maturation is completed, the mother cell lyses and the spore is released into the environment. The different layers of the spore are indicated: coat, OM (outer spore membrane), cortex and IM (inner spore membrane). **(B)** Schematic of germination. Germination is initiated by the binding of germinants (orange spheres) to germinant receptors (GR). This is followed by a series of events which lead to the release of DPA, hydrolysis of spore peptidoglycan, and rehydration of the spore. During germination, the spore transitions from phase-bright to phase-dark. Phase-contrast images of a mature phase-bright spore (left) and a germinated phase-dark spore (right) are shown next to the schematic. Scale bar, 1µM.

Under favorable conditions, spores germinate and resume vegetative growth (14–16). Spores sense the environment through the detection of specific germinant molecules that are recognized by receptors in the inner spore membrane (Fig. 1B). Germinants vary between species, but are typically small metabolites such as amino acids, nucleotides, or sugars. In some cases, the presence of a single germinant is sufficient to induce germination. In other cases, multiple germinants need to be present simultaneously. Germinant receptors are very sensitive to these molecules, and germination can sometimes be initiated by germinants at micromolar concentrations (17, 18). The binding of germinants to their receptors triggers the release of cations from the spore, which is followed by the release of DPA, hydrolysis of the spore cortex peptidoglycan, and rehydration of the spore core. After germination, the spore loses its heat resistance and appears phase-dark under phase-contrast microscopy (Fig. 1B).

Although the use of a common nutrient as a germinant would appear to be a sound strategy for detecting a favorable environment in which to emerge from dormancy, this strategy also carries with it the risk of untimely germination, particularly during spore formation. Sporulation is an elaborate developmental program that requires the synthesis of more than 500 different proteins (19), many in large quantities. Some of these proteins are components of the spore, and others participate in the intricate spatial and temporal regulation that enables the program to progress through its series of stages. Extensive protein synthesis of course requires an abundance of amino acids, and some of these amino acids are also potent germinants. Alanine or valine, for example, can each trigger germination of *B. subtilis* spores (20), so the very amino acids that the sporulating *B. subtilis* needs to carry the program through to completion could also subvert the process by initiating inappropriate germination. Were this to happen, it would be extremely disadvantageous for the population. Provoked by the threat of imminent starvation to embark upon a resource-intensive developmental program leading to the production of a resilient spore, these cells would instead produce only a newly germinated vegetative cell that would likely face an even bleaker environment than that which prompted its progenitor to commit to sporulation. In order to prevent the loss of their investment in this way, sporulating bacteria must be able to reduce the levels or suppress the activity of any germinants that are present during sporulation.

Here we show how sporulating *B. subtilis* accomplishes this. We find that catabolism of the germinant amino acids alanine and valine prevents the untimely germination of *B. subtilis* spores in their sporulation media, and that spores produced by mutants that are unable to catabolize these amino acids germinate prematurely due to their accumulation in the culture medium. We infer that clearance of germinant amino acids is required to prevent futile sporulation and germination. This finding has important physiological, ecological and evolutionary implications.

## RESULTS

### Strains that lack alanine dehydrogenase produce spores that lose their heat resistance during extended incubation in their spent sporulation medium

Germination of *B. subtilis* spores is typically triggered by small nutrient molecules such as alanine or valine, which could then be catabolized after germination to provide carbon, nitrogen, and energy for growth (21). We were interested in the contribution of the intrinsic metabolic potential of dormant *B. subtilis* spores to enabling growth after germination, so we wanted to make use of a germinant that was not toxic, metabolically inert, and could not double as a nutrient. No such germinant has been described for *B. subtilis*, but we reasoned that we could turn alanine into the inert germinant that we needed by simply preventing its catabolism. Alanine dehydrogenase (Ald) catalyzes the first reaction in alanine catabolism, and is required for growth on alanine as the sole carbon or nitrogen source (22–25). The enzyme couples the oxidative deamination of alanine to the reduction of NAD^+^ to produce pyruvate and ammonium (22, 26). But our simple strategy was complicated by the prior observation that Ald is required for efficient spore production in *B. subtilis* (24). We wondered whether this would still be true if sporulation was carried out in a medium that contained no alanine to be catabolized. We therefore assessed the sporulation efficiency of an Ald^−^ mutant in DSM, the conventional complex sporulation medium, which contains alanine both as the free amino acid and incorporated into peptides (27, 28), and also in S7_50_, a defined minimal medium that contains no alanine, but only glucose, glutamate and ammonium as carbon and nitrogen sources (29, 30). We determined the titers of heat-resistant spores for the wild-type (Ald^+^) and Ald^−^ strains in each medium, and we did so over several days because we anticipated different growth rates and sporulation efficiencies in these two different media.

Spore titers of the Ald^+^ strain peaked at greater than 10^8^ spores/mL in both media, at 24 h (day 1) in the complex DSM (Fig. 2A) and typically at 48 h in the minimal S7_50_ (Fig. 2B; Fig. S1). These titers remained constant in both media as the culture was sampled over the next few days (Fig. 2A and B; Fig. S1). The Ald^−^ strain, in contrast, showed unexpected spore titer profiles in both media. In DSM, spore titers were almost two orders of magnitude lower than those of the Ald^+^ strain at 24 h, in close agreement with the prior observation (24). But we were surprised to see the spore titer gradually fall off during subsequent days by about an additional order of magnitude, such that at seven days, the titer was roughly three orders of magnitude below that of the Ald^+^ strain (Fig. 2A). Sporulation of the Ald^−^ strain appeared to be more efficient in S7_50_ than in DSM. The titers in S7_50_ were only slightly lower than those of the Ald^+^ strain during the first day or two (Fig. 2B; Fig. S1). However, allowing for some variability early in the time course (Fig. S1), spore titers dropped by about two and a half orders of magnitude over the following days, to approximately three orders of magnitude below those of the Ald^+^ strain. It appeared that the spores of the Ald^−^ strain had become compromised merely through prolonged incubation in the medium in which they were sporulated.

**Figure 2.**
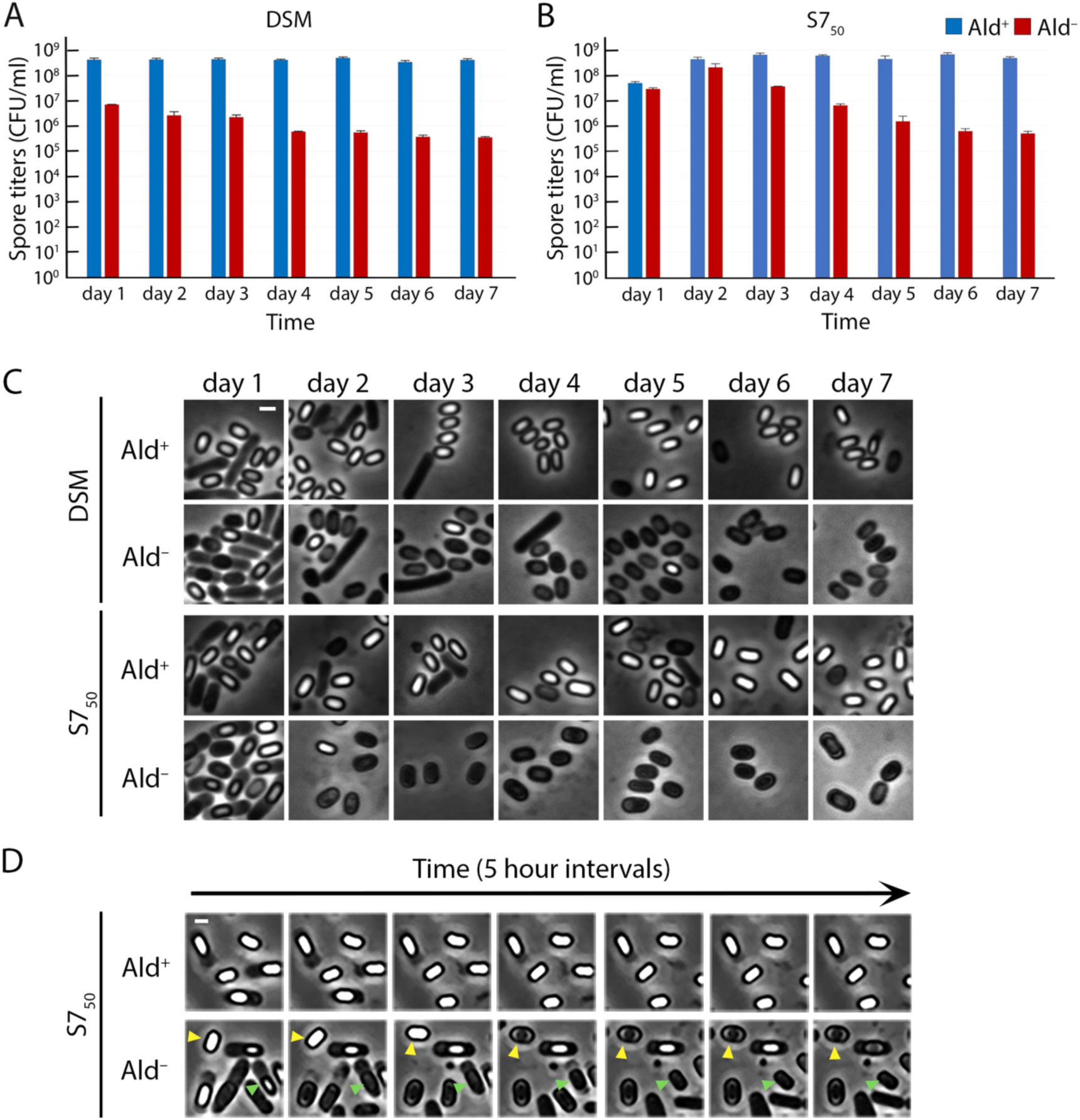
Ald^−^ spores germinate prematurely in their sporulation medium. Heat-kill spore titers of wild type (Ald^+^, blue bars) and Ald^−^ (red bars) strains in **(A)** DSM and **(B)** S7_50_. Data represent the average and standard deviation of three independent cultures. **(C)** Phase-contrast microscopy of the Ald^+^ and Ald^−^ strains in DSM and S7_50_ media. Samples were imaged every day for seven days. Scale bar, 1 µm. **(D)** Images from timelapse phase-contrast microscopy of sporulating cells from Ald^+^ and Ald^−^ cultures (Movies S1 and S2). One field is shown at 5-hour intervals for the first 30 hours of the experiment. Within this field are a spore that transitions to phase-dark while it is still within the mother cell (green arrowheads) and a free spore that turns phase-dark (yellow arrowheads). Scale bar, 1 µm.

### Spores in Ald^−^ cultures germinate prematurely

We sampled Ald^+^ and Ald^−^ cultures at different days and visualized spores using phase-contrast microscopy. Mature heat-resistant spores have a low water content and appear bright in the phase-contrast microscope. Heat-sensitive spores have a higher water content and are phase-dark (Fig. 1B). The Ald^+^ culture contained largely free phase-bright spores at 24 h in DSM and at 48 h in S7_50_. Free phase-bright spores persisted in both cultures throughout the rest of the incubation (Fig. 2C). This was as expected, given the typical and consistent heat-kill spore titers we recorded for the wild-type strain (Fig. 2A and B).

The Ald^−^ culture contained fewer free phase-bright spores than did the Ald^+^ culture at 24 h in DSM (Fig. 2C). Some spores were phase-dark and still within the mother cell; either they had never become phase-bright, or they had transitioned from phase-bright back to phase-dark while in the mother cell. In S7_50_ at 24 h, the Ald^−^ culture contained many spores within the mother cell, much like the Ald^+^ culture at the same timepoint. However, the Ald^−^ spores tended to be less bright than those of the Ald^+^ culture, suggesting that the spores of the Ald^−^ cultures were maturing slowly or were unable to mature fully inside their mother cells. At later timepoints, both DSM and S7_50_ Ald^−^cultures contained mostly free phase-dark spores, indicating that the phase-bright spores that had been present at earlier timepoints had lost their phase-brightness in the spent media.

We were able to capture the transition from phase-bright to phase-dark in real time with time-lapse microscopy. We examined the S7_50_ cultures at 24 h, when the Ald^+^ and Ald^−^ cultures contained similar amounts of developing spores, most still within the mother cells (Fig. 2C). The cells were loaded into a microfluidic chip and incubated with their culture supernatants for an additional 56 hours (Fig. 2D; Movies S1 and S2). During the course of the timelapse, the majority of the Ald^+^ spores remained phase-bright (Fig. 2D; Movie S1). We did observe that a small fraction of Ald^+^ spores (5.3%; n = 359) transitioned from phase-bright to phase-dark (Movie S1), which is consistent with what has been reported (31). During the same time, however, 77.1% (n = 576) of the phase-bright Ald^−^ spores transitioned to phase-dark (Movie S2). We observed this transition with phase-bright spores that were still within the mother cell (Fig. 2D, green arrowheads), frequently in conjunction with mother cell lysis, and also with free phase-bright spores (Fig. 2D, yellow arrowheads).

During germination, spores are rehydrated and lose their phase-brightness and heat-resistance. The spores in the Ald^−^ cultures appeared to be germinating in their spent medium. This would explain the transition from phase-bright to phase dark in the Ald^−^ cultures and the decrease in the number of heat-resistant spores (Fig. 2). The inner membrane of *B. subtilis* spores contains three receptors, GerA, GerB and GerK, that mediate the initiation of germination in response to particular molecules in the environment (32). Either alanine or valine can on its own trigger germination via GerA (20), but no single germinant has been found to trigger germination via GerB or GerK. Instead, GerB and GerK initiate germination jointly in response to a mixture of asparagine, glucose, fructose and potassium ions (AGFK) (33, 34). In order to determine whether the sporulation defects observed in Ald^−^ cultures were due to premature germination, we inactivated each receptor and determined the spore titers over the course of several days. Inactivation of GerB or GerK did not affect the spore titer profiles of Ald^+^ and Ald^−^ cultures in either DSM or in S7_50_ (Fig. S2). However, inactivation of GerA eliminated completely the spore titer defect of the Ald^−^ strain in both media (Fig. 3A and B). Timelapse phase-contrast microscopy of S7_50_ cultures confirmed that in the absence of the GerA receptor, Ald^−^ spores were phase-bright and did not transition to phase-dark (Fig. 3C; Fig. S3; Movies S1 to S4). GerA has been previously implicated in the premature germination of abnormal spores (31). We therefore infer that premature germination is responsible for the sporulation defect of an Ald^−^ mutant, and that this premature germination is triggered through the GerA receptor.

**Figure 3.**
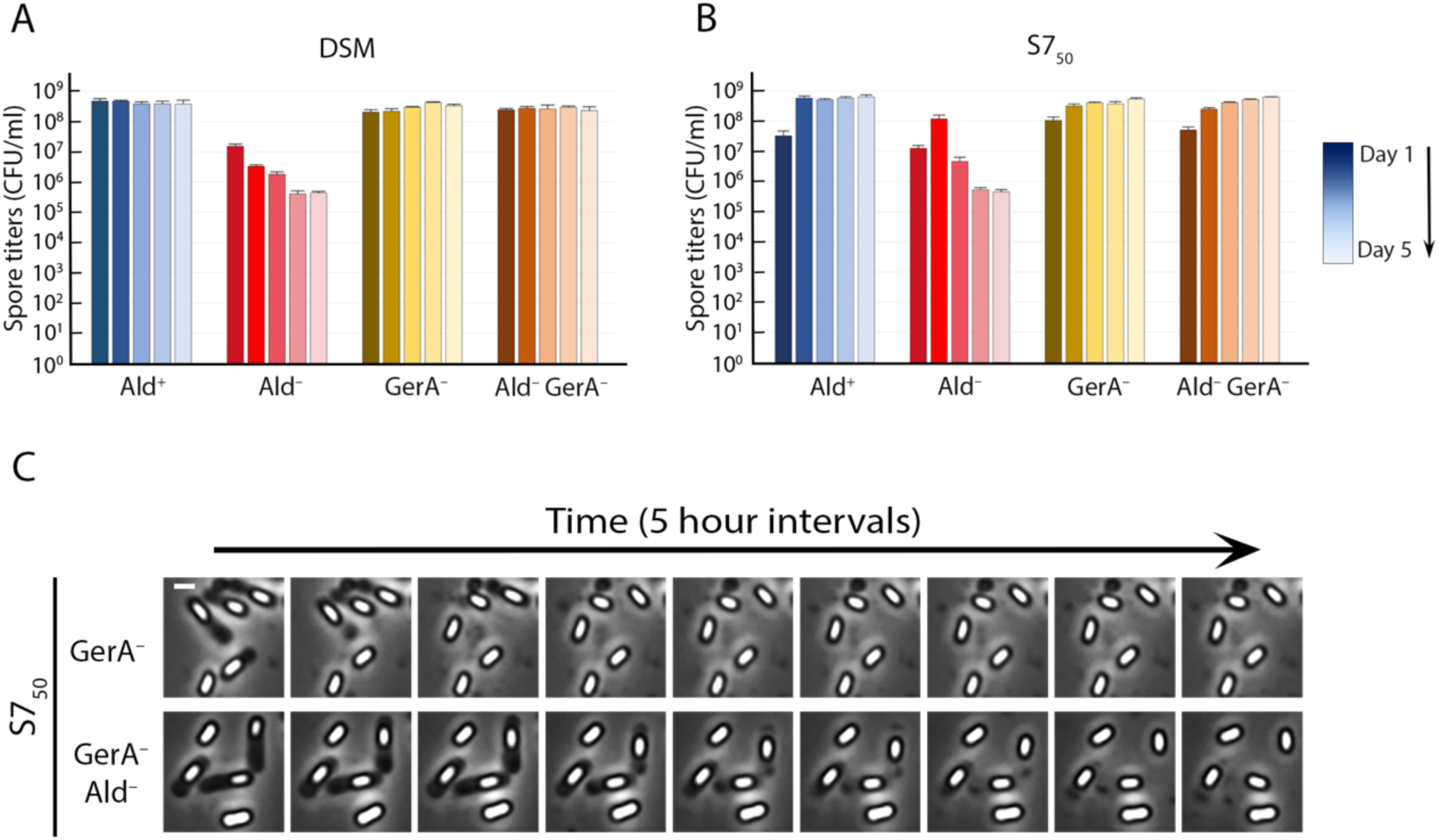
Elimination of GerA prevents premature germination of Ald^−^ spores. Heat-kill spore titres in **(A)** DSM and **(B)** S7_50_ of isogenic Ald^+^ and Ald^−^ strains containing or lacking GerA (GerA^−^). Blue bars, Ald^+^ GerA^+^; red bars, Ald^−^ GerA^+^; brown to yellow bars, Ald^+^ GerA^−^; brown to beige bars, Ald^−^ GerA^−^. The color gradient tracks the time course, from Day 1 (darkest shade) to Day 5 (lightest shade). Spore titers were determined daily; the data are the average and standard deviation of three independent cultures. **(C)** Images from timelapse phase-contrast microscopy of sporulating Ald^+^ (upper row) and Ald^−^ (lower row) strains in a GerA^−^ background (from Movies S3 and S4). One field is shown at 5-hour intervals for the first 40 hours of the experiment. Scale bar, 1 µm. See Figure S3 for images from timelapses of isogenic Ald^+^ and Ald^−^ strains in GerA^+^ and GerA^−^ backgrounds.

### Accumulation of alanine in the medium is responsible for the premature germination of Ald^−^ spores

The GerA receptor triggers germination in response to alanine or valine. It seemed likely that the sporulation defects in Ald^−^ cultures were caused by the accumulation of unmetabolized alanine. We therefore determined the free amino acid content of spent DSM and S7_50_ from Ald^+^ and Ald^−^ cultures. Fresh DSM contains beef infusion and an enzymatic digest of animal protein (27), so it was unsurprising to find many free amino acids at high µM and even mM concentrations. Alanine was present at 1.3 mM, one of the highest concentrations (Table 1, Table S1). But after 48 h of growth of the Ald^+^ strain in DSM, by which point the culture consisted of mostly free phase-bright spores, the amount of alanine had dropped dramatically, by more than 95% (Table 1, Table S1). The cells had consumed all of the free alanine in the medium, and presumably also the alanine that was released from hydrolysis of the DSM peptides. The Ald^−^ culture looked very different. After 48 h, the spent DSM contained nearly 3 mM alanine, more than twice the concentration in the fresh medium (Table 1). The Ald^−^ strain had failed to consume the alanine, and peptide hydrolysis served merely to add more alanine to the unconsumed pool. The situation was similar when S7_50_, which contains no alanine, was used for sporulation. Alanine was undetectable in fresh S7_50_, as expected, and also in the spent S7_50_ when the Ald^+^ strain was grown for 48 h (Table 1). But after the Ald^−^ strain was grown for 48 h, alanine was present at about 400 μM, owing presumably to lysis and to leakage from the cells (Table 1). The Ald^−^ strain failed to scavenge this alanine.

**Table 1.**
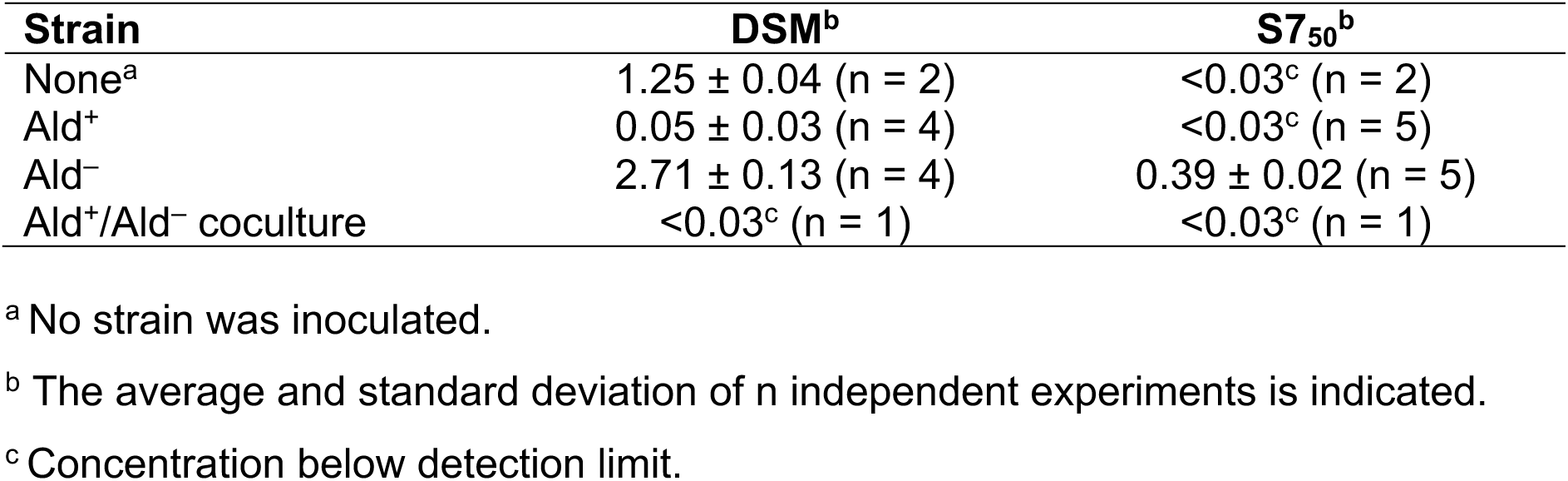
Alanine concentration (mM) after growth of the indicated strain for 48 h.

| Strain | DSM<sup>b</sup> | S7<sub>50</sub><sup>b</sup> |
| --- | --- | --- |
| None<sup>a</sup> | 1.25 ± 0.04 (n = 2) | <0.03<sup>c</sup> (n = 2) |
| Ald<sup>+</sup> | 0.05 ± 0.03 (n = 4) | <0.03<sup>c</sup> (n = 5) |
| Ald<sup>-</sup> | 2.71 ± 0.13 (n = 4) | 0.39 ± 0.02 (n = 5) |
| Ald<sup>+</sup>/Ald<sup>-</sup> coculture | <0.03<sup>c</sup> (n = 1) | <0.03<sup>c</sup> (n = 1) |

<sup>a</sup> No strain was inoculated.
<sup>b</sup> The average and standard deviation of n independent experiments is indicated.
<sup>c</sup> Concentration below detection limit.

In both DSM and S7_50_, the concentration of alanine in the spent medium of the Ald^−^ culture was well above the 60 μM required to achieve 50% of the maximum germination rate (18). Attainment of this concentration would be expected to be more unpredictable in S7_50_ than in DSM because of the stochastic nature of the lysis and leakage events that supply alanine to the medium, and this in turn likely accounts for the variability that we observed early in the S7_50_ time courses (see above and Fig. 2B, Fig. S1). When we supplemented S7_50_ with alanine so as to match its concentration in DSM, the drop in spore titers was always apparent at day 1, and the profile came to resemble that which we observed with DSM (Fig. S4).

To confirm that it was the alanine that was responsible for triggering germination, we grew and sporulated the Ald^−^ and Ald^+^ strains together in a single DSM or S7_50_ coculture. On the basis of the amino acid analysis, we expected the Ald^+^ strain to catabolize the alanine that the Ald^−^ strain was failing to consume (Fig. 4A). If premature germination was triggered by alanine, it should not occur in the coculture. The strains were first modified to express genes for distinct fluorophores from a constitutive variant of the P*_spank_* promoter (35), so that they could be distinguished under the microscope and on agar plates (Fig. 4A and B). The spore titer profile of the fluorescently-labeled Ald^+^ and Ald^−^ strains growing singly matched those of the isogenic unlabeled strains: the Ald^+^ spore titers remained stable, and the Ald^−^ spore titers dropped off over time (Fig. 4C and D). Coculturing of the Ald^−^ and Ald^+^ strains, either in DSM or in S7_50_, eliminated the drop off in spore titers that we had observed when the Ald^−^ strain was cultured singly (Fig. 4C and D). The Ald^−^ spores remained phase-bright in coculture even though they transitioned prematurely to phase-dark when the strain was grown singly (Fig. 4B). Amino acid quantification confirmed that no alanine was detectable in the spent medium of the coculture (Table 1).

**Figure 4.**
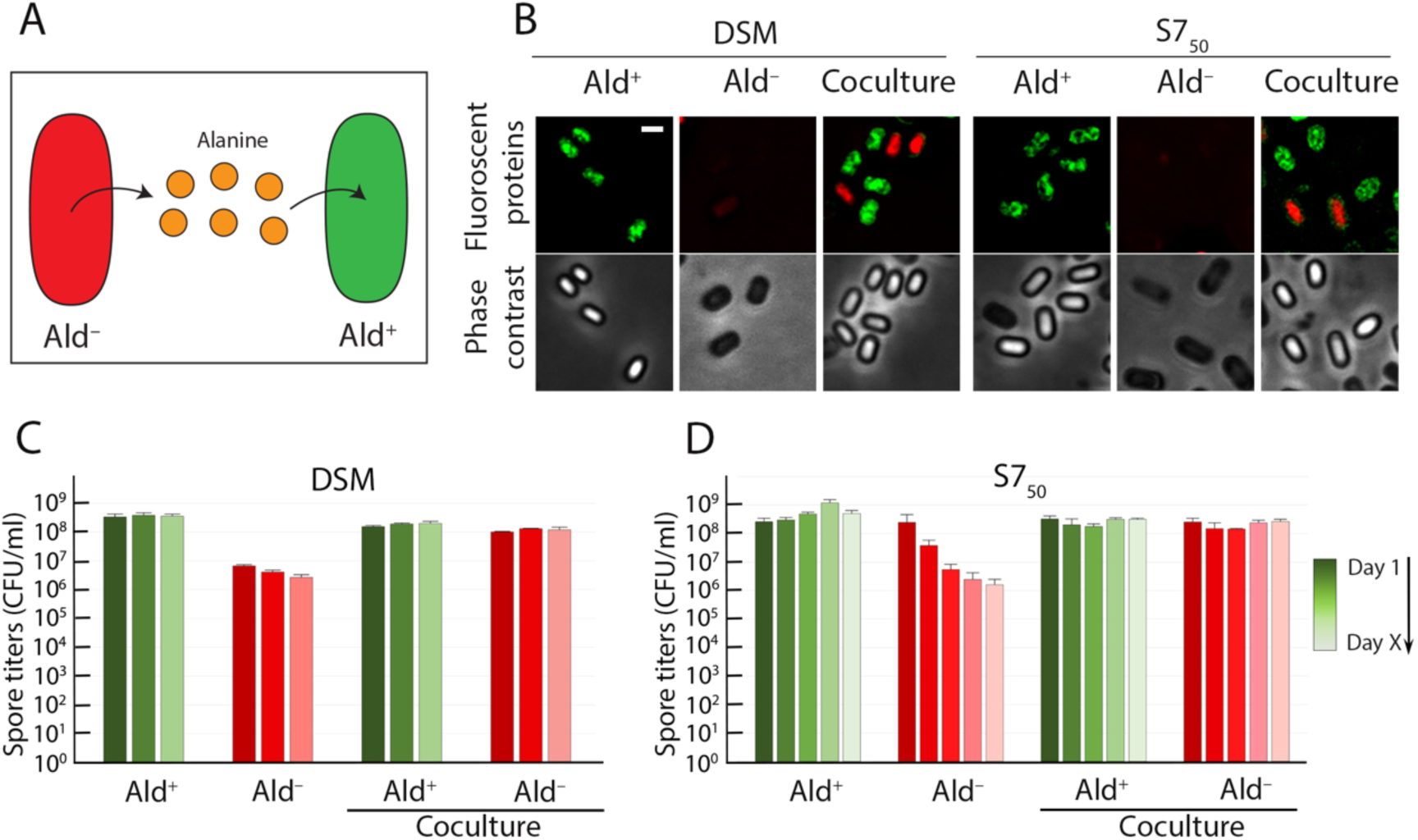
Alanine scavenging prevents the premature germination of Ald^−^ spores. **(A)** In a population containing both Ald^+^ and Ald^−^ cells, the unmetabolized alanine released by or from the Ald^−^ cells (red), will be scavenged by the Ald^+^ cells (green) and removed from the medium. **(B)** Fluorescence (top row) and phase-contrast (bottom row) images of Ald^+^ and Ald^−^ strains (monoculture) and their coculture after 48 hours of incubation in DSM or S7_50_ media. The Ald^+^ cells produce mNeonGreen (green) and the Ald^−^ cells produce Tomato (red) constitutively. These fluorescent proteins can be detected in phase-bright spores but not in phase-dark spores (see Methods). Scale bar, 1 µm. Heat-kill spore titers of Ald^+^ and Ald^−^ strains in monocultures and cocultures in **(C)** DSM and **(D)** S7_50_. The color gradient tracks the time course, from Day 1 (darkest shade) to Day X (lightest shade). X = 3 for DSM; X = 5 for S7_50_. Graph shows the averages and standard deviations of three independent cultures.

### The falling spore titers reflect only premature germination, not failure to outgrow

Alanine can function as a germinant during germination and then as a nutrient during the outgrowth that follows. The microscopy data demonstrate that the defect that is associated with the buildup of alanine in the Ald^−^ strain is due only to alanine in its capacity as a germinant (Fig. 2C and D; Fig. S3). But the spore titer measurements that support these data report on both germination and outgrowth (*e.g.* Fig. 2A and B). In order to determine whether we had to account for outgrowth when evaluating our spore titers, we needed to determine what contribution Ald makes to outgrowth. If Ald contributes little or is actually dispensable during outgrowth on our germination medium (LB), then the diminished spore titers must be due only to the premature germination defect. We therefore produced Ald^−^ spores and compared their outgrowth on LB to that of wild-type Ald^+^ spores.

We engineered a strain that contained Ald during vegetative growth and sporulation, but produced spores that lacked Ald and that were therefore unable to catabolize alanine during outgrowth. To do this, we targeted Ald for *in vivo* degradation using a strategy that we have used before to eliminate specific proteins from the mother cell or forespore during spore formation (36–39). We modified the *ald* gene so that Ald was produced with a C-terminal ssrA tag (35), and we produced the *E. coli* protease-targeting factor SspB^Ec^ in the forespore late in the course of sporulation (Fig. 5A). This yields a strain that sporulates as Ald^+^ but produces spores that are Ald^−^ (Fig. 5B). This (Ald^+^) strain sporulated normally with no premature germination (Fig. S5, upper row). The resulting (Ald^−^) spores also outgrew normally in LB (Fig. S5C, middle row) and the spore titers matched those of the wild-type strain (Fig. 5C and D; Fig. S5D and E). The spores truly lacked Ald—they germinated but did not outgrow when alanine was the only carbon source (Fig. S5C, lower row). Inducing the degradation of the ssrA-tagged Ald in vegetative cells prevented them from growing on alanine, again confirming that Ald could be eliminated from the cell with this strategy (Fig. S5A and B). In short, Ald^−^ spores outgrew as well as wild-type spores in LB, so Ald makes no significant contribution during outgrowth, and the falling spore titers of the Ald^−^ strain therefore reflected only the premature germination defect. That the spore titers of an Ald^−^ strain were equivalent to those of the wild-type either when GerA was inactivated (Fig. 3 A and B) or in coculture with an Ald^+^ strain (Fig. 4C and D) underscored this conclusion.

**Figure 5.**
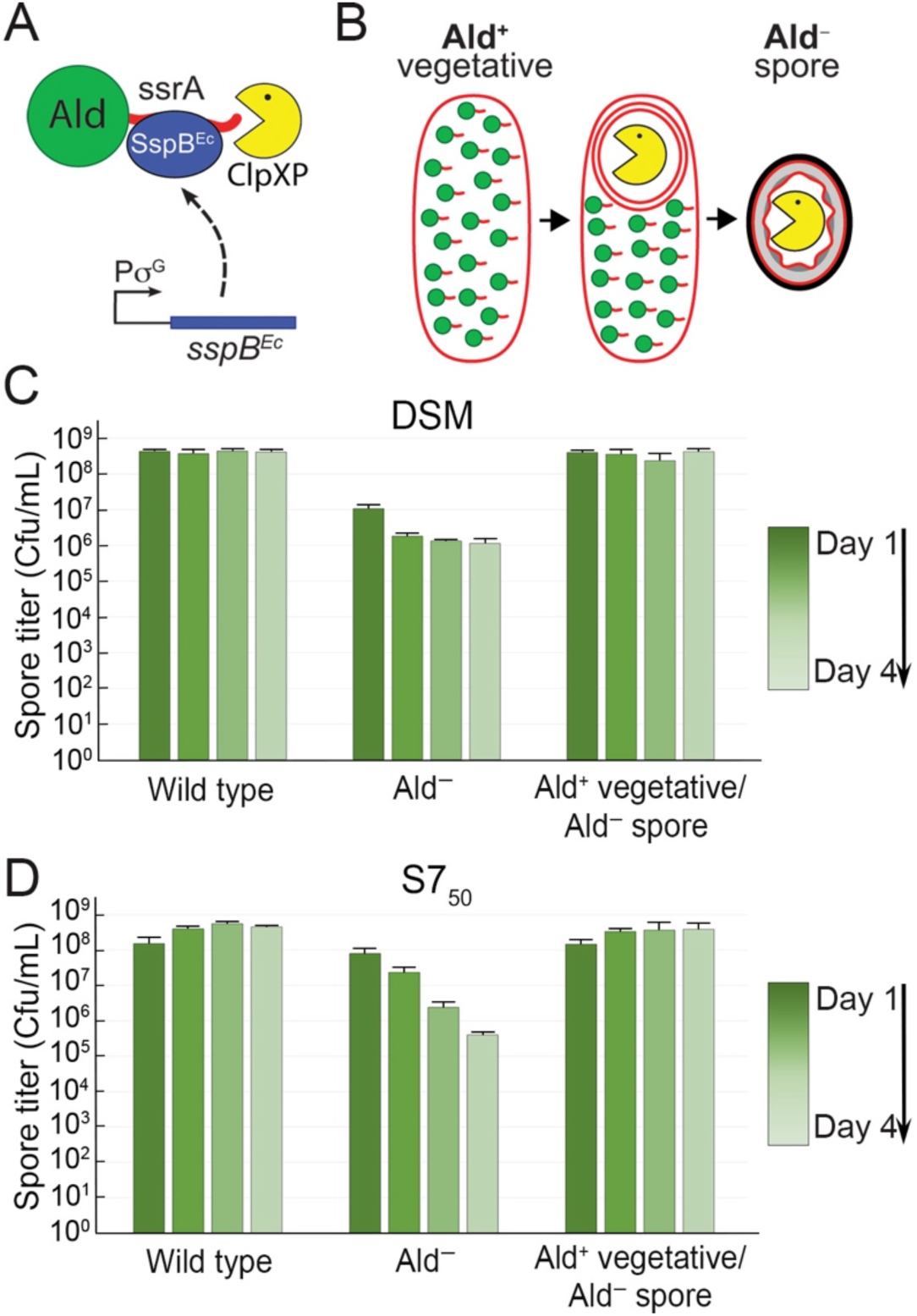
Alanine dehydrogenase is dispensable during outgrowth in rich medium. **(A)** Schematic of targeted Ald degradation. Ald (green sphere) is tagged with ssrA (red tail). The adaptor protein SspB^Ec^ (blue oval) binds to the ssrA-tag and delivers Ald-ssrA to the ClpXP protease (yellow pacman) for degradation. The *sspB*^Ec^ gene is expressed from a σ^G^-dependent promoter that is activated in the forespore after engulfment is completed. **(B)** Ald-ssrA (green sphere with red tail) is not degraded during vegetative growth, so vegetative cells are Ald^+^. Ald-ssrA is degraded only during sporulation, only in the forespore (yellow pacman), and only after engulfment is complete, leading to the production of Ald^−^ spores (See also Fig. S5). **(C, D)** Heat-kill spore titer profiles in DSM **(C)** or S7_50_ **(D)** of the wild-type strain, the Ald^−^ strain, and of the strain in which Ald-ssrA is degraded late in the forespore during sporulation, leading to the production of Ald^−^ spores (Ald^+^ vegetative/Ald^−^ spore). The color gradient tracks the time course, from Day 1 (darkest) to Day 4 (lightest). Spore titers were determined daily and the data are the average and standard deviation of three independent cultures.

### A mutant that cannot catabolize valine germinates in its spent medium

When Ald is not present, the uncatabolized alanine triggers premature germination of *B. subtilis* spores, either during their development or shortly after they mature. Do defects in the catabolism of other amino acids also lead to premature germination? We went on to assess the sporulation proficiency of *B. subtilis* mutants that lack one of these other catabolic enzymes: AnsA (asparagine; (40)), AnsB (aspartic acid; (40)), Bcd (valine, leucine, and isoleucine [branched-chain amino acids]; (41)), HutH (histidine; (42)), SdaA (serine; (43)), and Tdh (threonine; (44)). The catabolic reactions that these enzymes catalyze are diagrammed in Fig. 6A.

**Figure 6.**
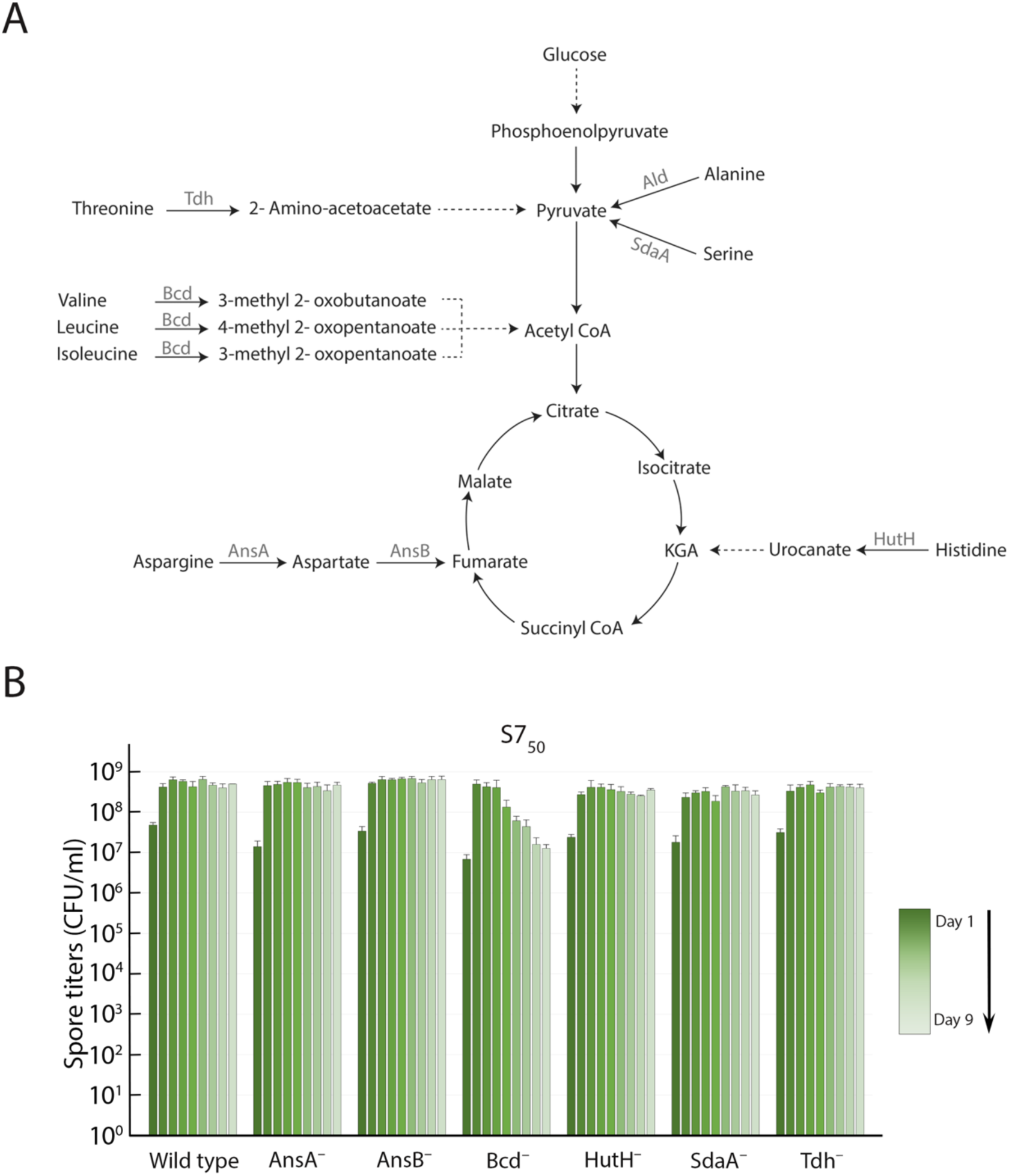
Spore titer profiles of mutants unable to catabolize particular amino acids. **(A)** Catabolic pathways for the amino acids examined in the study. The first enzyme in each pathway, the reaction that it catalyzes (solid arrow), and the product generated are shown; subsequent catabolic steps are represented as dashed arrows. **(B)** Heat-kill spore titers in S7_50_ medium of the wild-type culture and of cultures of mutants that lack the indicated enzymes. The green gradient tracks the time course, from Day 1 (darkest) to Day 9 (lightest). Spore titers were determined daily; the data are the average and standard deviation of three independent cultures.

We determined the titer of heat-resistant spores every 24 hours over nine days (Fig. 6B; Fig. S6), as above. The AsnA^−^, AsnB^−^, HutH^−^, SdaA^−^, and Tdh^−^ strains achieved and maintained spore titers that were similar to those of the wild-type. Only the Bcd^−^ strain had a profile resembling that of the Ald^−^ strain. In both DSM and S7_50_, the spore titers peaked at 48 h and then fell off during the following days. The peak titer was delayed by about one day with respect to that of the wild-type strain in both media, most likely owing to the slower growth of the Bcd^−^ strain (not shown). In DSM, the peak was followed by a modest decline (Fig. S6). In S7_50_, the peak spore titer persisted through the fourth day and then fell off sharply over several days (Fig. 6B).

Microscopy of samples taken from S7_50_ cultures of the Bcd^−^ strain at different days revealed that phase-bright Bcd^−^ spores turned phase-dark over time, suggesting that they were germinating prematurely in their spent medium (Fig. 7A). Like alanine, valine can trigger spore germination through the GerA receptor. Because Bcd initiates the catabolism of valine, we suspected that valine was accumulating in its absence and triggering premature spore germination in much the same way as did alanine in the Ald^−^ strain. Valine was indeed present in the spent medium of the Bcd^−^ strain in DSM at 0.87 mM and in S7_50_ at 0.18 mM, but was undetectable in the spent medium of wild-type (Bcd^+^) cultures (Table S2). Leucine and isoleucine, the other Bcd substrates, were present as well (not shown), but these amino acids are not known to trigger germination of *B. subtilis* spores. Although a valine concentration of 3.7 mM was reported to be required to achieve 50% of the maximum germination rate (18), we found that at 0.5 mM valine, 40% of wild-type spores germinate in 5 h, and that some germination was apparent at 0.2 mM as well (not shown).

**Figure 7.**
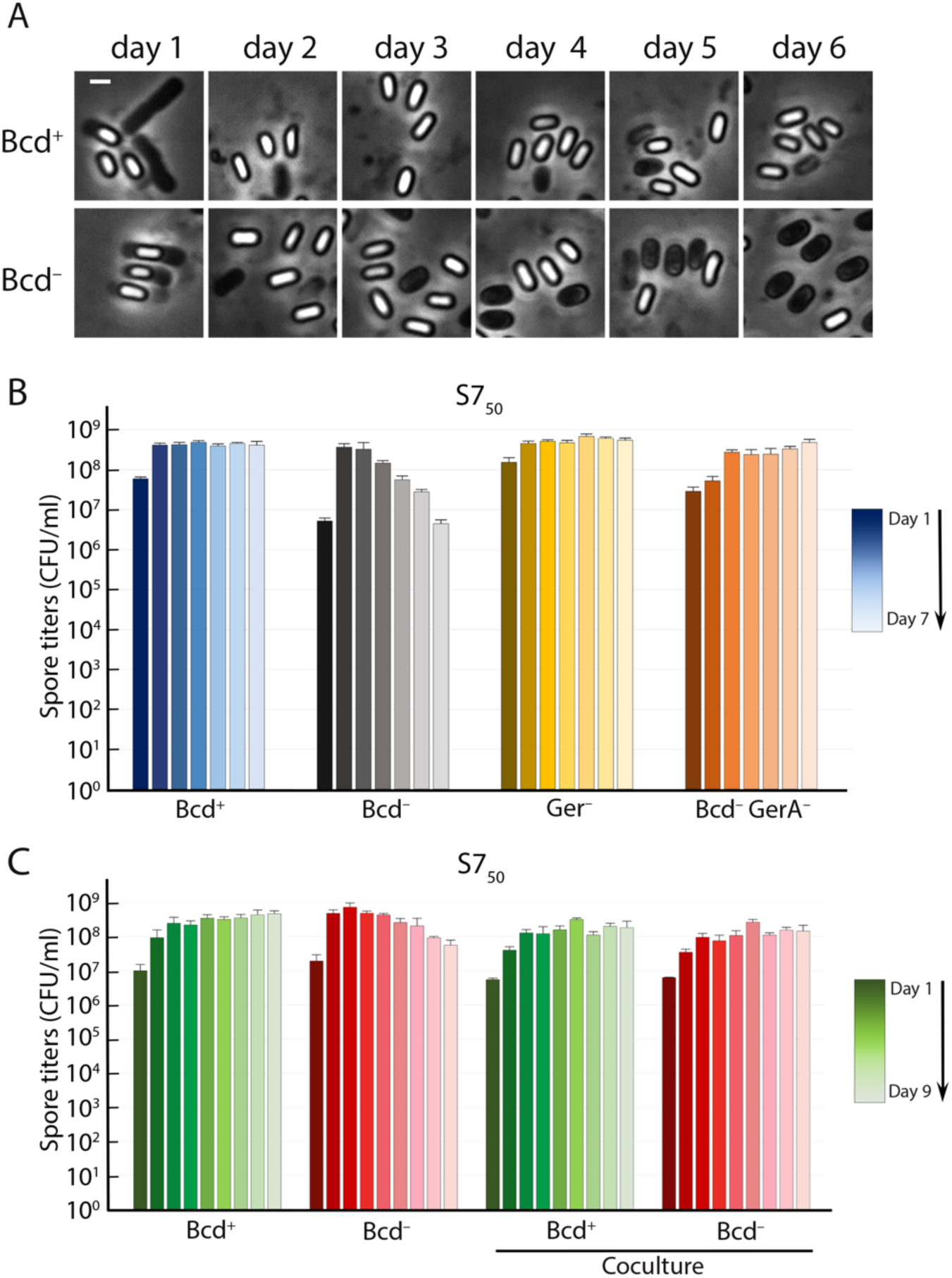
Bcd^−^ spores germinate prematurely in their spent medium. **(A)** Phase-contrast microscopy of Bcd^+^ and Bcd^−^ strains in S7_50_ medium. Images were captured every day for six days. Scale bar, 1 µm. **(B)** Heat-kill spore titers in S7_50_ of isogenic Bcd^+^ and Bcd^−^ strains containing or lacking GerA (GerA^−^). Blue bars, Bcd^+^ GerA^+^; black to grey bars, Bcd^−^ GerA^+^; brown to yellow bars, Bcd^+^ GerA^−^; brown to beige bars, Bcd^−^ GerA^−^. The color gradient tracks the time course, from Day 1 (darkest) to Day 7 (lightest). Spore titers were determined daily; the data are the average and standard deviation of three independent cultures. **(C)** Heat-kill spore titers of Bcd^+^ and Bcd^−^ strains in monocultures and cocultures in S7_50_. The color gradient tracks the time course, from Day 1 (darkest) to Day 9 (lightest). Spore titers were determined daily; the data are the average and standard deviation of three independent cultures.

The inactivation of the GerA receptor in the Bcd^−^ strain completely suppressed the spore titer defects (Fig. 7B). No suppression was observed when GerB or GerK, which do not respond to valine as a germinant, were inactivated (Fig. S7). In addition, coculturing of the Bcd^−^ strain with a Bcd^+^ strain that could catabolize valine prevented the germination of the Bcd^−^ spores in the spent media and eliminated the defect completely (Fig. 7C). The profile therefore matched that which we had observed for the Ald^−^ strain with respect to alanine, indicating that the same mechanism was in play here.

## DISCUSSION

We have shown that *B. subtilis* prevents futile sporulation and germination by clearing the amino acids alanine and valine before the developing spore can respond to them as germinants. Clearance is accomplished by Ald and Bcd, which initiate the catabolism of alanine and valine, respectively. In the absence of these enzymes, the germinant amino acids accumulate in the sporulation medium and trigger premature germination of the newly formed spores.

Our results have prompted us to revisit the explanation of the role of Ald in sporulation that emerged from previous studies (24). Using transposon mutagenesis, Sandman and collaborators isolated insertions that impaired sporulation (45). One of the insertions was in a locus initially called *spoVN*, and caused a blockage at late stages of sporulation that was characterized by the failure of the spores to accumulate DPA (45). It was later shown that *spoVN* was actually *ald*, and it was proposed that alanine catabolism was necessary to provide energy for sporulation (24). We have demonstrated however that sporulation can proceed normally in the absence of alanine catabolism, as Ald^−^ mutants form mature spores at wild-type levels so long as the alanine that accumulates is removed or the GerA receptor is inactivated. We have also shown that Ald is not required for outgrowth in rich medium. We conclude that the lack of DPA accumulation in Ald^−^ spores and the spore titer defects of Ald^−^ strains are not due to a specific requirement for alanine catabolism to fuel sporulation or outgrowth, but are instead manifestations of premature germination.

It follows from our findings that metabolism and germinant selection are tightly linked in *B. subtilis*. This linkage has physiological, ecological, and evolutionary implications. The physiological implication is that metabolism must be configured so that germinants are preferentially cleared prior to or during spore formation. For amino acids such as alanine and valine, clearance requires their catabolic amino acid dehydrogenases, Ald in the case of alanine, and Bcd in the case of valine. The *B. subtilis* Ald is well-adapted to this task, with a *K*_m_ of 70 μM and a *k*_cat_/*K*_m_ of nearly 2 x 10^6^ M^-1^s^-1^ (46). Clearance could in theory begin once the commitment to sporulation has been made, or it could be ongoing during vegetative growth and an integral part of metabolism. Regulational studies point to the latter scenario, and to one in which clearance is so critical that the familiar hierarchies of catabolite repression are dispensed with. Alanine and valine induce the transcription of their catabolic dehydrogenases even in the presence of preferred carbon sources such as glucose (24, 41, 47–49). Ald levels peak at the onset of sporulation, even in the absence of exogenous alanine (21, 24), and this could ensure that alanine is cleared by the early stages of the program. Valine, leucine, and isoleucine activate the transcriptional regulator CodY (50), which represses the expression of genes required for the synthesis of branched-chain amino acids, contributing to the maintenance of low levels of valine inside the cells (51). Although other amino acids are cleared equally efficiently, some accumulate during growth in wild-type cultures (Table S1). When we grew *B. subtilis* in a defined medium that contained each amino acid at 3 mM, about half of the amino acids were cleared after 2 days of growth, including alanine and valine. But others, such as methionine and lysine, were present at high concentrations in the spent medium (Table S1). Lysine was present in spent DSM and also in spent S7_50_, which contains no lysine to begin with (Table S1). Thus, the clearance priorities for the amino acids differ, with the germinant amino acids having high or perhaps highest priority. One might expect similar behavior from other endospore-forming bacteria for which alanine can serve as a germinant. Spores from *B. amyloliquefaciens* germinate in response to alanine (52). We found this to be true as well of *B. mojavensis*, *B. spizezenii*, and *B. clausi* (not shown). All of these endospore-formers reduced the alanine concentration in the growth medium to undetectable or near undetectable levels in the course of 48 h. Non-spore-forming bacteria gave a varied profile, where some species consumed all of the alanine and others left much of it in the medium (Table S3).

Our findings raise the question of how spore formers reconcile the need to eliminate germinant amino acids with the need to preserve them for protein biosynthesis and, in the case of alanine, peptidoglycan biosynthesis as well. The answer to this question remains unclear at this point. However, interconversion between L-alanine and the D-alanine that is required for peptidoglycan biosynthesis, as mediated by alanine racemase, would seem to provide a solution in the case of alanine. Unlike L-alanine, D-alanine is not a germinant, and it actually inhibits the binding of L-alanine to GerA (53). Stores of D-alanine could help to prevent premature germination and at the same time supply L-alanine to the cell as needed. Other species of *Bacillus*, notably *Bacillus anthracis*, may use precisely this strategy to modulate L-alanine levels during sporulation. Several different species of endospore-forming bacteria encode an alanine racemase that is produced specifically during sporulation (54–56). In *B. anthracis*, the absence of the sporulation alanine racemase causes the spore to germinate inside the mother cell during development (57), which strongly suggests that alanine racemase performs the same L-alanine clearing function for *B. anthracis* that Ald performs for *B. subtilis*. It is possible that *B. subtilis* uses its sporulation-specific alanine racemase in the same way, but if so, the enzyme makes only a minor contribution, as *B. subtilis* lacking its sporulation-specific alanine racemase does not show premature germination (31, 58).

The ecological implications of our results are hard to assess because we understand very little about *B. subtilis* beyond the conventional laboratory monoculture. Perhaps a *B. subtilis* that lacked the ability to remove alanine or valine would not be at a competitive disadvantage in the soil because microbes in the vicinity would readily scavenge these essential amino acids. One wonders though whether relying on competing microbes for germinant clearance would be a sound strategy. The range of nutrients that microbes take up varies widely, even among strains of the same species (59), so the extent to which a particular germinant would be depleted would depend upon the constitution of the surrounding microbial community. And if sporulation occurs within the context of *B. subtils* biofilms, as is likely the case, there would be few other species present to clear the germinant. *B. subtilis* cells that are committed to sporulation first produce factors that kill non-sporulating sibling cells and other microbes (60, 61), and the resulting cell lysis would likely lead to the release of germinants. These would need to be promptly cleared, and this would be most likely to happen if the sporulating cells did this themselves, if they could rely upon an intrinsic mechanism to clear germinants prior to spore formation.

It is well understood that endospore-forming bacteria carry distinct germinant receptors that respond to specific germinants or sets of germinants (32). Not nearly as well understood are the selective forces that determine which molecules function as germinants. The present study reveals a major constraint on germinant selection: the germinant must be metabolically compatible with the bacterium—the cell must be able to clear the germinants before the receptor is present and functioning in the envelope of the developing spore. Clearance would typically require catabolism of the germinant, but it could involve incorporation of the germinant into macromolecules, or in the special case of alanine, racemization to the D-enantiomer. Because different bacterial species and sometimes even different strains of the same species have different metabolite consumption profiles, in particular with respect to amino acids (59), a suitable germinant for one endospore-former may not be a suitable germinant for another. In the case of *B. subtilis*, for example, lysine is always present in spent media, and therefore could not function as a germinant (Table S1). An endospore-forming bacterium that acquired a receptor for a molecule that it could not clear would accrue a significant fitness cost from premature germination, and there would be strong selective pressure for a modification to the receptor or to its catabolic priorities. The evolution of germinant receptors and the evolution of metabolism must therefore be inextricably connected.

Many germinant receptors respond to combinations of molecules, rather than to single molecules (32). For example, the *B. subtilis* GerB and GerK receptors require D-glucose, D-fructose, L-asparagine, and K^+^ for activation (20, 33). If one of the four is absent, the others are unable to together trigger germination, so clearing only one would be sufficient to prevent premature germination. The requirement for multiple germinants to trigger a single germinant receptor has been understood as enabling the dormant spore to assess the availability of more than just one nutrient before committing to emerging from dormancy. This requirement may now also be understood as having evolved as a safeguard against premature germination during sporulation.

## MATERIALS AND METHODS

### Strains and plasmids

The strains used in this study are listed in Table S4. Strains constructed for this study are listed along with schematics of their construction. Strains previously constructed, already in our laboratory collection, or obtained from stock centers or other investigators are listed along with their sources. Gene disruptions were transferred from the BKK and BKE knockout strain collection (62) into *Bacillus subtilis* PY79 by transformation. The resulting strains were verified by PCR in which flanking primers were used to amplify the region containing the disruption, or a flanking primer and one matching the introduced antibiotic cassette were used for amplification. Oligonucleotide primer sequences used for constructions and verifications are listed in Table S6. Plasmids are listed in Table S5, followed by descriptions of their construction.

### Culture media and growth conditions

*B. subtilis* PY79 and derivatives were routinely grown on LB agar plates at 30°C. Liquid cultures were grown at 37°C with shaking at 220 rpm. Spore titres were determined in two different liquid media: (i) DSM, a complex medium containing protein fragments, peptides, and free amino acids [(27, 63); Table S1]. DIFCO nutrient broth was obtained from Beckton-Dickinson (Catalog #234000). (ii) S7_50_, a defined minimal medium, was as described, but with thiamine at 3 μM (29, 30). Glucose and glutamate were each at 20 mM; ammonium sulfate was at 10 mM (as per the recipe). The medium used to determine the amino acid catabolic preferences of *B. subtilis* PY79 and other spore formers contained all 20 amino acids at 3 mM as sole carbon source (Table S1). Cysteine and methionine were supplied from 50 mM stocks that were kept at room temperature for one week or less. The other amino acids were supplied from 50 mM and 12.5 mM pools that were stored in aliquots at −80°C. The medium was based on S7_50_ and contained the S7_50_ salts and metals at their prescribed concentrations (as above, with 3 μM thiamine), but no glucose, and no glutamate other than the 3 mM in the amino acid mixture. *B. subtilis* PY79 sporulated efficiently in this medium (not shown). The agar plates shown in Fig. S5 contained 10 mM alanine with or without 1% xylose.

### Quantification of heat-resistant spores

Culture volumes for sporulation were 2 or 10 mL, depending upon the number of samplings required for the experiment. Cultures were incubated at 37°C and shaken at 220 rpm. The smaller, 2 mL cultures were grown in 10 mL tubes, and incubated in an air shaker. The larger, 10 mL cultures were grown in 100 mL baffled flasks and incubated in a shaking water bath. Every 24 h, 50 µL was abstracted and heated at 80°C for 20 min, and then serially diluted tenfold in T-base buffer (0.2% (NH_4_)_2_SO_4_, 1.4% K_2_HPO_4_, 0.6% KH_2_PO_4_, 0.1% Na_3_Citrate_・_H_2_O). Five μL of each dilution was spotted onto LB agar plates, the plates were incubated overnight at 30°C, and spore titers were calculated from the number of colonies that arose.

### Phase-contrast and fluorescence microscopy

Cultures were imaged on pads of 1.2% agarose in T-base buffer. Ten µL of the culture was deposited directly onto the pad, the liquid was allowed to dry, and the pad covered with a standard #1.5 glass coverslip. A DeltaVision Ultra fluorescence deconvolution microscope equipped with a PCO Edge sCMOS camera was used for image acquisition. For phase-contrast microscopy, light transmission was set to 100% and exposure time to 0.1 s. Tomato and mNeonGreen were visualized with the appropriate filters: excitation 575/25 and emission 625/45 for Tomato, and excitation 475/28 and emission 525/48 for mNeonGreen, with light transmission at 30% and an exposure time of 0.1s. Fluorescent images were deconvolved with SoftWoRx. Images for the manuscript were prepared with ImageJ2 image processing software (version 2.9.0 / 1.53v).

### Timelapse phase-contrast microscopy

A CellASIC® ONIX2 microfluidic system with a Manifold Basic for pressure control was used for timelapse microscopy with B04A CellASIC® ONIX plates. S7_50_ cultures were prepared by diluting logarithmic phase cultures (OD_600_ ∼ 0.6) to an OD_600_ of 0.1, and then incubating for 24 h at 37°C with shaking at 220 rpm. One mL of the culture was centrifuged at 9,400 g for 5 min to pellet the cells, and the supernatant was recovered. After the cells were loaded into the microfluidics plate, the spent media was flowed through at a pressure of 2.5 PSI for 8 h, at the end of which time the flow was stopped. Phase-contrast images were collected every 5 min for 56 h on our DeltaVision Ultra microscope (above). Light transmission was set to 100%, and the exposure time was 0.1 s.

### Purification of spores

Spores were purified with a scaled-down version of the published protocol (64). Cultures grown for 48 hours were centrifuged at 4500 g for 15 min at 4°C, and the pellet was washed 3 times with 5 mL of cold (4°C) sterile water. The supernatant was removed, the pellet was resuspended in the volume remaining, and 600 uL 3 M potassium phosphate buffer (1.24 M monobasic/1.76 M dibasic) and 900 uL 50% PEG 4000 were added. The mixture was vortexed vigorously for 2 to 5 min, and then spun at 6000 g for 10 min at 4°C. The spore-containing organic phase (top) was transferred to a new tube, diluted with 1 mL of cold sterile water, and the suspension was spun at 6000 g for 5 min at 4°C. The spore pellet was washed twice with 1 mL cold water, resuspended in 500 uL cold water, and a sample examined with phase-contrast microscopy. If the preparation contained phase-dark spores, a further purification by HistoDenz gradient was carried out (65). The impure spore pellet was resuspended in 20% HistoDenz, the suspension was layered onto 1.5 mL of 50% HistoDenz, and centrifuged at 8000 g for 10 min at 4°C. The pellet of phase-bright spores was washed three times in cold water with the same centrifugation protocol. The pellet was resuspended in fresh sterile water, and stored at 4°C. Sample purity was evaluated with phase-contrast microscopy.

### Phase contrast microscopy of purified spores

Spores were imaged on pads of 1.2% agarose in LB or minimal media (S7_50_ base + 1 mM alanine). Ten µL of the purified spore preparation was deposited directly onto the pad, the liquid was allowed to dry, and the pad was covered with a standard glass coverslip and sealed to the slide with petroleum jelly to avoid dehydration during incubation. The slides were incubated at 30°C for 6 hours (LB) or 24 hours (minimal media). Phase contrast microscopy was as described above.

### Amino-acid scavenging experiment

Strains were marked with fluorescent proteins in order to differentiate them in coculture. The wild-type strain was labelled with mNeonGreen (66) and the Ald^−^ and Bcd^−^ strains were labelled with Tomato (red) (67). Both proteins were produced from genes under the control of the constitutive Pc promoter (35), and were detectable during sporulation and in the mature spores. Fluorescence was not detected in germinated spores in spent media (Figure 4B), likely because the fluorescent proteins were degraded during germination. Precultures of each strain in DSM or S7_50_ were grown into logarithmic phase (OD_600_ ∼ 0.6) and diluted into 5 ml of fresh media to an OD_600_ of 0.01. These cultures were grown to an OD_600_ of approximately 0.1, and each was either mixed with its partner for the experiment in a 1:1 ratio, or continued as a single culture, in either case with a final volume of 2 mL. The cultures were shaken at 220 rpm at 37°C for 3 to 5 days, depending upon the media. Spore titers were determined daily as described above.

### Amino acid quantification analysis

Logarithmic phase cultures (OD_600_ ∼ 0.6), either DSM or S7_50_, were diluted to an OD_600_ of 0.1 in 10 mL of media and grown at 37°C with shaking at 220 rpm. Culture supernatants were typically harvested at 48 h, with the exception of the supernatants of Bcd^−^ cultures and their respective wild-type (Bcd^+^) controls, whose supernatants were harvested at 120 h of incubation. Cultures were centrifuged at 4,500 g for 10 min, and the supernatants filtered through a 0.2 µm cellulose acetate membrane syringe filter. HPLC (High Performance Liquid Chromatography) amino acid analysis of the supernatants was carried out by Sartorius Xell GmbH (Göttingen, Germany). The experimental blanks were also submitted for analysis.

## ACKNOWLEDGEMENTS

We are grateful to all of the members of Evolutionary Cell Biology research group and to the Microbial Population Biology department of the MPI for Evolutionary Biology for useful discussion and comments during the development of this project. We acknowledge the National BioResource Project (NIG, Japan): *B. subtilis* for shipping the BKE collection to us and the *Bacillus* Genetic Stock Center (Columbus, Ohio) for other spore-forming species. We are grateful to Daniel Unterweger and Olga Volger for the non-spore forming species used in this study, and for assistance with the experiments with pathogenic bacteria that were performed in a biosafety level 2 facility. This work was supported by the European Research Council (ERC) Starting Grant 853383 and by the Max-Planck-Gesellschaft. The authors declare no conflicts of interest.

