## Supplementary Figures and Tables for "Catabolism of germinant amino acids is required to prevent premature spore germination in *Bacillus subtilis*"

- Figures S1 to S7
- Tables S1 to S6
- Legend for Movies S1 to S4

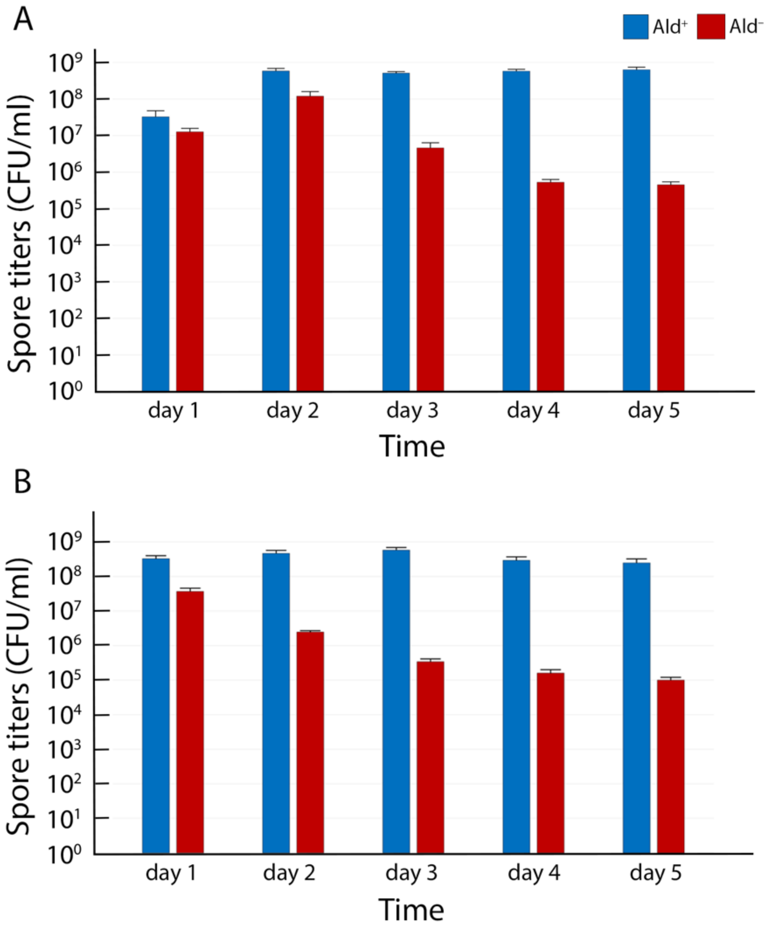


**Figure S1. Variability in the $S7_{50}$ spore titers at early time points.** Heat-kill spore titers of $Ald^+$ (blue bars) and $Ald^-$ (red bars) strains in $S7_{50}$ medium. Panels **(A)** and **(B)** show results from two independent experiments, each performed in triplicate. In the first experiment **(A)**, the titers of both the $Ald^+$ and $Ald^-$ strains increased at day 2 compared to day 1. In the second experiment **(B)**, the titers peaked at day 1 for both strains. In both experiments, the $Ald^-$ titers dropped during the following days, reaching values that were about three orders of magnitude below those of the $Ald^+$. The data in each graph are the average and standard deviation of three independent cultures.

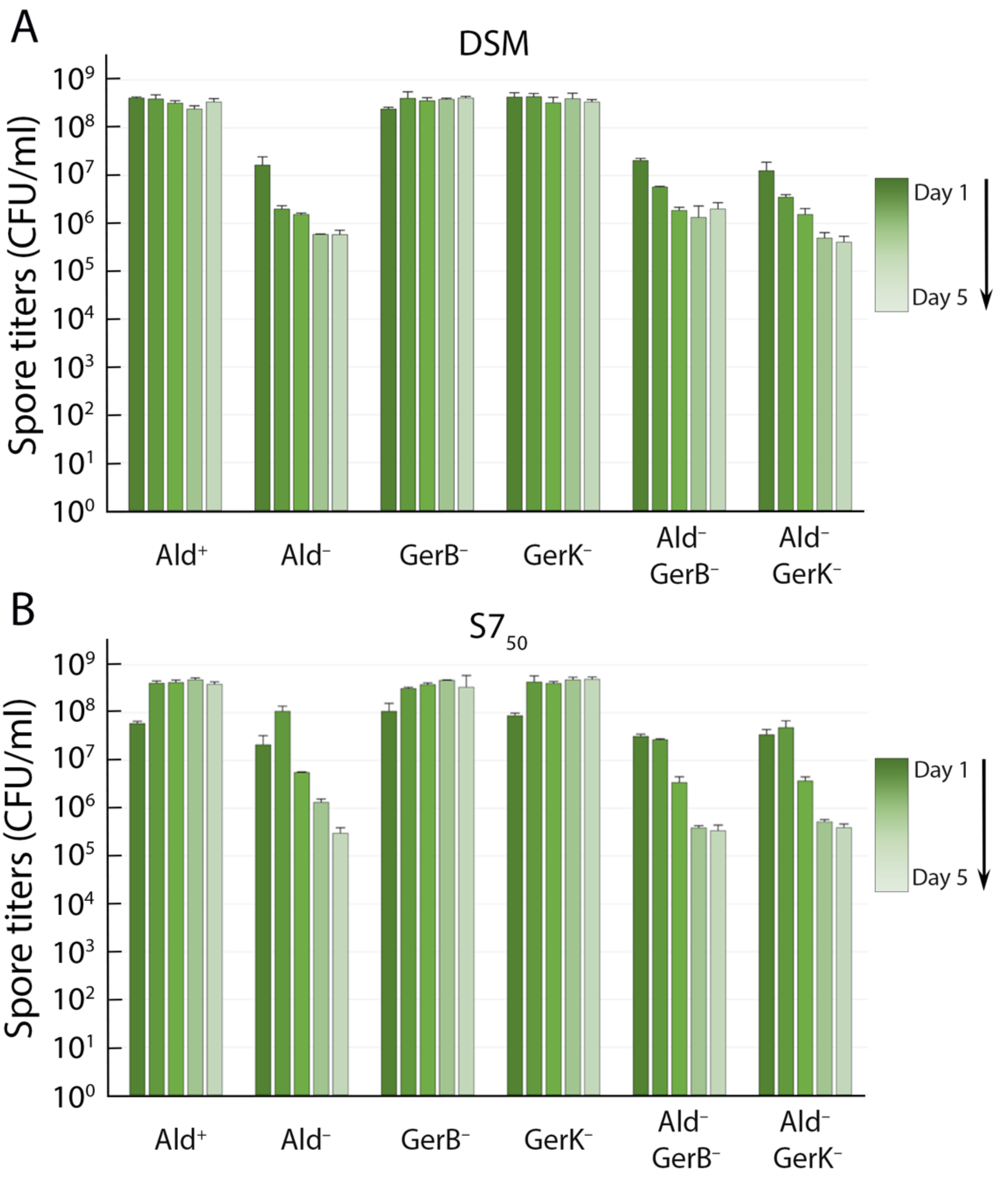


**Figure S2. Elimination of GerB or GerK does not prevent premature germination of Ald⁻ spores.** Heat-kill spore titers of isogenic Ald⁺ and Ald⁻ strains containing or lacking the GerB (GerB⁻) or GerK (GerK⁻) receptors in **(A)** DSM and **(B)** $S7_{50}$. The color gradient tracks the time course, from Day 1 (darkest) to Day 5 (lightest). Spore titers were determined daily; the data are the average and standard deviation of three independent cultures.

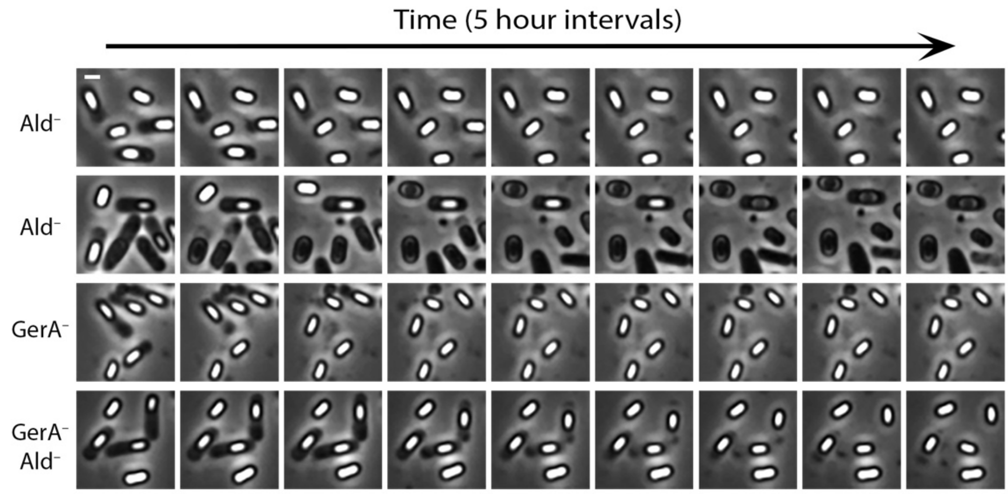


**Figure S3. Overview of timelapse microscopy.** Images from timelapse phase-contrast microscopy of sporulating cells from $Ald^{+}$ (first row) $Ald^{-}$ (second row), $GerA^{-}$ (third row) and $Ald^{-}$ $GerA^{-}$ cultures (Movies S1 to S4). Images taken at 5-hour intervals are shown. The brightness and contrast of every picture has been adjusted using the same settings, to facilitate the visual comparison of spore brightness. Scale bar, 1 µm.

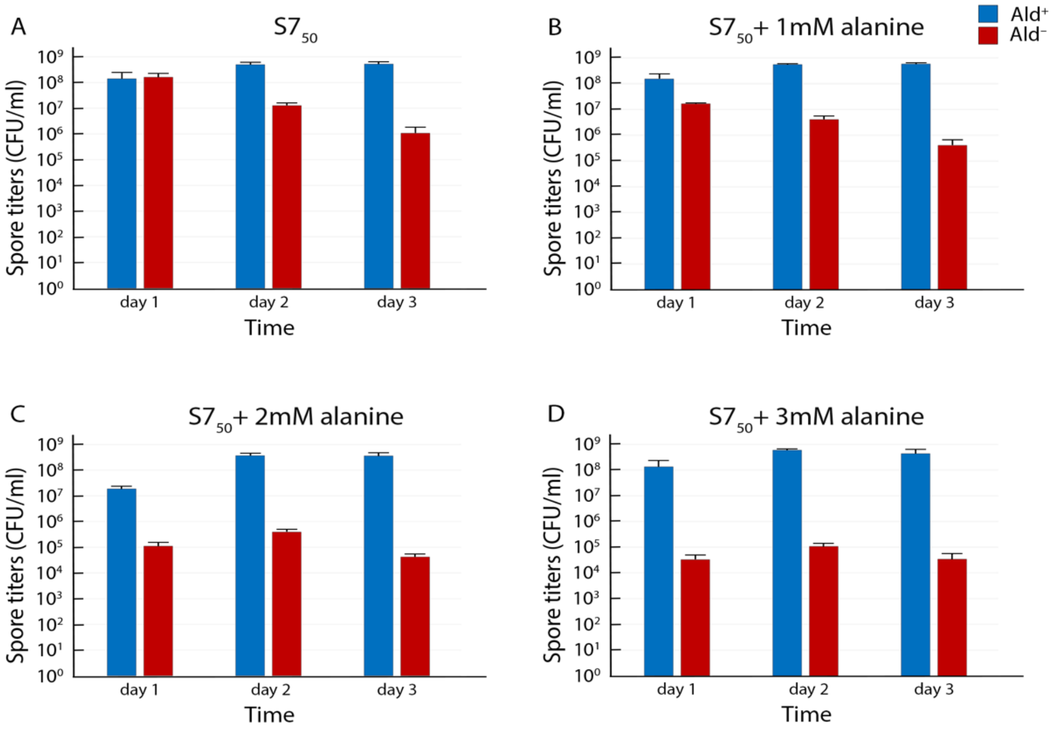


**Figure S4. Ald⁻ spore titers in S7$_{50}$ supplemented with alanine.** Heat-kill spore titers of Ald⁺ (blue bars) and Ald⁻ (red bars) strains in **(A)** S7$_{50}$ medium, **(B)** S7$_{50}$ +1mM alanine **(C)** S7$_{50}$ +2mM alanine, and **(D)** S7$_{50}$ +3mM alanine. The data in each graph are the average and standard deviation of three independent cultures.

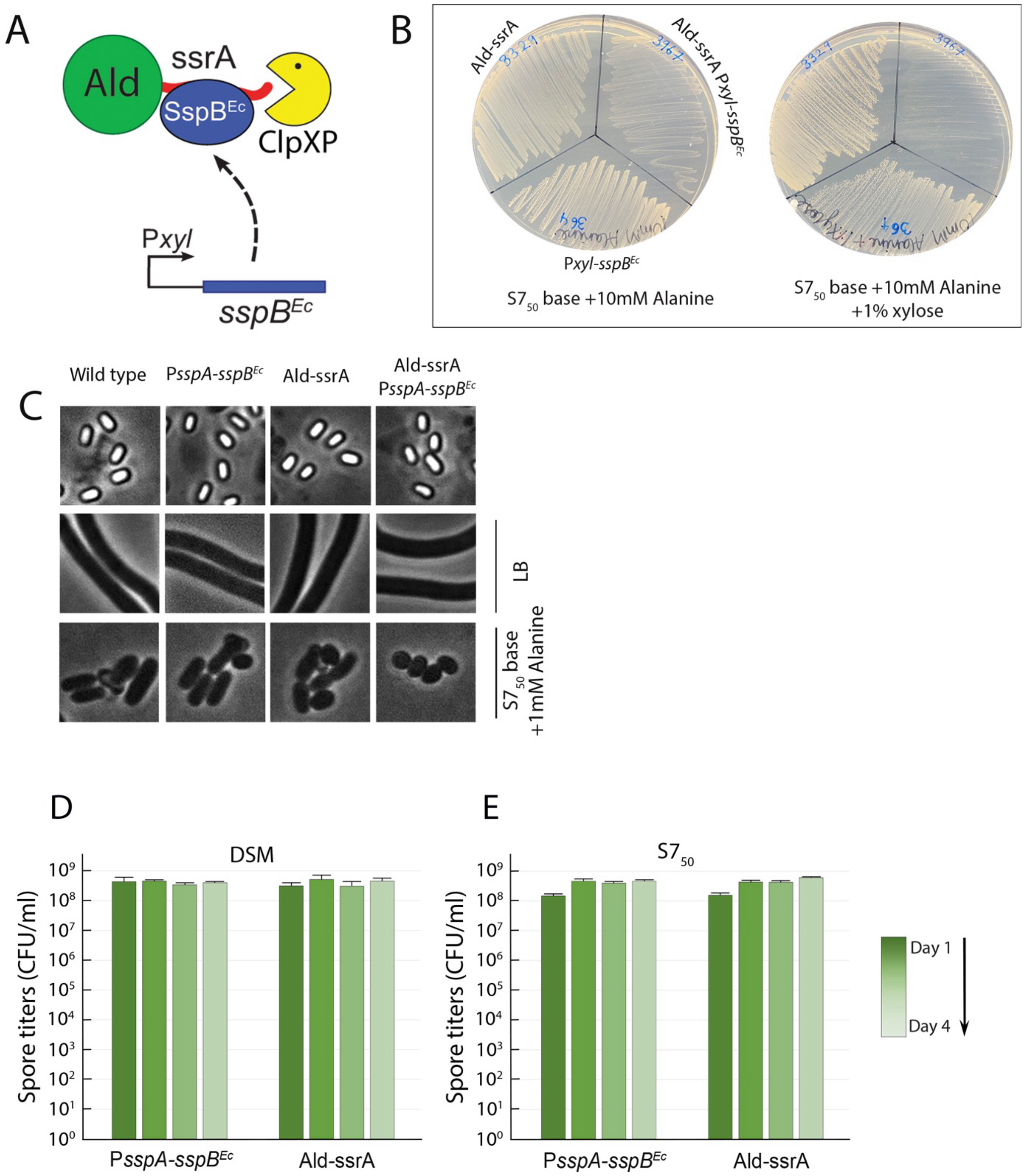


**Figure S5. Alanine dehydrogenase tagged with ssrA is efficiently degraded. (A)** Schematic of xylose-inducible Ald degradation. Ald (green sphere) is tagged with ssrA (red tail). The adaptor protein SspB$^{Ec}$ (blue oval) binds to the ssrA-tag and delivers Ald-ssrA to the ClpXP protease (yellow pacman) for degradation. The *sspB$^{Ec}$* gene is expressed from the xylose-inducible promoter P*xyl*. **(B)** Ald-ssrA is efficiently degraded when *sspB$^{Ec}$* is expressed in the cell. Left, S7$_{50}$ minimal medium base with 10 mM alanine as sole carbon source; right, the same, but with 1% xylose. Strains are: (*i*) Ald-ssrA, in which Ald is produced with the ssrA tag, but the strain contains no *sspB*$^{Ec}$ gene; (*ii*) P*xyl-sspB*$^{Ec}$, which contains *sspB*$^{Ec}$ under the control of P*xyl*, but Ald is not tagged; and (*iii*) Ald-ssrA P*xyl-sspB*$^{Ec}$, which produces the tagged Ald, and *sspB*$^{Ec}$ is under P*xyl*. This strain fails to grow on plates containing xylose, indicating that the cells cannot utilize alanine as a carbon source. (The *sspB*$^{Ec}$ gene disrupts the operon that is required for xylose catabolism, so alanine is the only potential carbon source). This result confirms that Ald-ssrA is efficiently degraded when SspB$^{Ec}$ is produced. **(C)** Degradation of Ald-ssrA in the forespore leads to the production of spores that cannot catabolize alanine. Phase-contrast microscopy images of spores of (*i*) a wild-type strain; (*ii*) P*sspA-sspB*$^{Ec}$,which contains *sspB*$^{Ec}$ under control of a forespore-specific promoter, but in which Ald is not tagged; (*iii*) Ald-ssrA, a strain in which Ald is produced with the ssrA tag, but contains no *sspB*$^{Ec}$ gene; and (*iv*) P*sspA-sspB*$^{Ec}$ Ald-ssrA, which produces Ald-ssrA and contains P*sspA-sspB*$^{Ec}$. Upper row, images of purified spores. All strains sporulated efficiently and produced stable phase-bright spores. Middle row, images taken after incubation of purified spores in LB agarose pads for 6 h. Chains of vegetative cells are observed from all strains, showing that the spores germinated and outgrew efficiently in LB. Bottom row, images taken after incubation of purified spores in agarose pads containing 1 mM alanine as sole carbon source for 24 hours. Spores from all the strains germinated efficiently, with some outgrowth of the wild-type, P*sspA-sspB*$^{Ec}$, and Ald-ssrA strains, but spores from the Ald-ssrA P*sspA-sspB*$^{Ec}$ strain, in which Ald-ssrA was degraded in the forespore, failed to outgrow in the presence of alanine as sole carbon source. No outgrowth was observed even after 48 hours (not shown). This result confirms that Ald-ssrA degradation in the forespore leads to the production of spores that are phenotypically Ald$^{-}$. **(D, E)** Heat-kill spore titer profiles in DSM **(D)** or $S7_{50}$ **(E)** of the P*sspA-sspB*$^{Ec}$, and Ald-ssrA strains from Panel C. The spore titers resemble those of the wild-type strain (Fig. 5C and D), demonstrating that producing SspB$^{Ec}$ in the forespore (with Ald untagged), or tagging Ald with ssrA (with no *sspB*$^{Ec}$ gene present) does not impact spore titers. The color gradient tracks the time course, from Day 1 (darkest) to Day 4 (lightest). Spore titers were determined daily and the data are the average and standard deviation of three independent cultures.

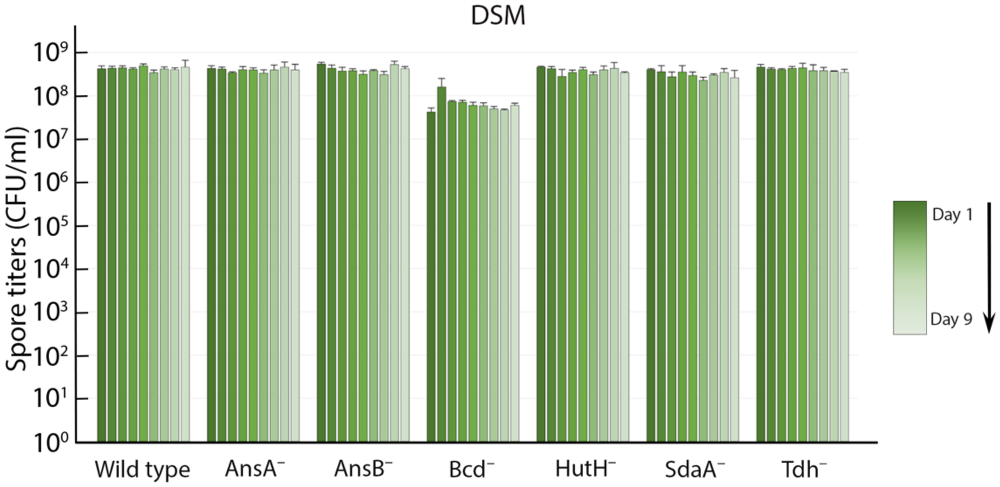


**Figure S6. DSM spore titer profiles of mutants unable to catabolize other amino acids.** Heat-kill spore titers in DSM media of the wild-type culture and of cultures of mutants that lack the indicated enzymes. The color gradient tracks the time course, from Day 1 (darkest) to Day 9 (lightest). Spore titers were determined daily; the data are the average and standard deviation of three independent cultures.

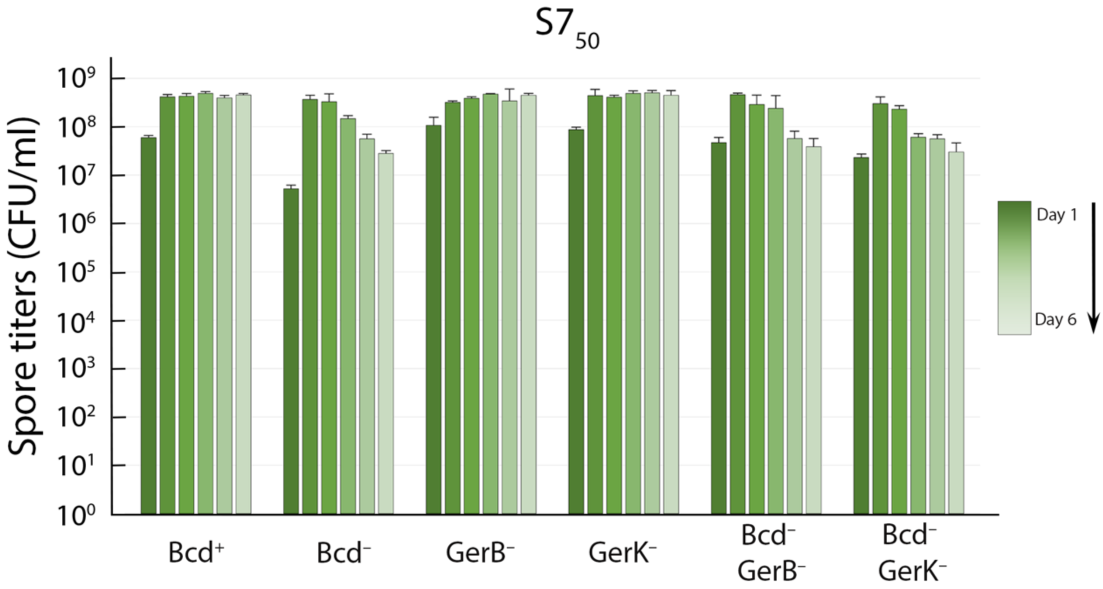


**Figure S7. Elimination of GerB or GerK does not prevent premature germination of Bcd⁻ spores.** Heat-kill spore titers in $S7_{50}$ of isogenic $Bcd^{+}$ and $Bcd^{-}$ strains containing or lacking the GerB ($GerB^{-}$) or GerK ($GerK^{-}$) receptors. The color gradient tracks the time course, from Day 1 (darkest) to Day 6 (lightest). Spore titers were determined daily; the data are the average and standard deviation of three independent cultures.

**Table S1.** Concentrations of free amino acids in sterile and spent media[a].

| | DSM | | $S7_{50}$ Equimolar[b] | |
|---|---|---|---|---|
| **Amino acids** | **Sterile (mM)** | **Spent(mM)** | **Sterile(mM)** | **Spent(mM)** |
| Alanine | 1.28 | <0.03[c] | 2.97 | <0.03[c] |
| Arginine | 1.69 | <0.03[c] | 2.95 | <0.03[c] |
| Asparagine | 0.23 | <0.03[c] | 2.83 | 0.14 |
| Aspartate | 0.54 | <0.03[c] | 2.95 | <0.03[c] |
| Cysteine[d] | <0.03[c] | <0.03[c] | <0.03[c] | <0.03[c] |
| Glutamine | <0.03[c] | <0.03[c] | 2.49 | <0.03[c] |
| Glutamate | <0.03[c] | 0.19 | 2.63 | 0.04 |
| Glycine | 1.13 | 0.35 | 3.09 | 0.30 |
| Histidine | <0.03[c] | <0.03[c] | 3.02 | <0.03[c] |
| Isoleucine | 0.40 | <0.03[c] | 3.22 | <0.03[c] |
| Leucine | 0.82 | <0.03[c] | 3.04 | <0.03[c] |
| Lysine | 1.00 | 0.96 | 3.03 | 2.60 |
| Methionine | 0.20 | 0.17 | 2.77 | 2.32 |
| Phenylalanine | 0.47 | 0.39 | 3.11 | 2.80 |
| Proline | 0.22 | <0.03[c] | 3.08 | 0.06 |
| Serine | 0.50 | 0.14 | 2.89 | <0.03[c] |
| Threonine | 0.34 | <0.03[c] | 2.92 | <0.03[c] |
| Tryptophan | 0.12 | <0.03[c] | 3.09 | 2.72 |
| Tyrosine | 0.32 | 0.18 | 3.06 | 2.74 |
| Valine | 0.59 | <0.03[c] | 2.90 | 0.05 |

[a]Spent media was recovered from the culture of an $Ald^+$ strain that was grown for 48 h at 37°C.

[b]$S7_{50}$ base ($S7_{50}$ salts and metals), to which was added each of the twenty amino acids to a concentration of 3 mM. See Materials and Methods.

[c]Concentration below detection limit.

[d]Cysteine was detected only in the form of cystine (not shown), owing to its oxidation in the stock solution or in the medium.

**Table S2.** Valine concentration (mM) after growth of the indicated strain for 120 h.

| **Strain** | **DSM**[b] | **$S7_{50}$**[b] |
|---|---|---|
| None[a] | 0.59 | <0.03[c] |
| Bcd$^+$ | <0.03[c] | <0.03[c] |
| Bcd$^-$ | 0.87 | 0.18 |

[a] Experimental blank

[b] Values from a single culture of each strain are reported.

[c] Concentration below detection limit.

**Table S3.** Alanine concentration (mM) after growth of the indicated strains for 48 h in $S7_{50}$ equimolar medium[a].

| Strain | Alanine concentration[c] |
|---|---|
| None[b] | 2.68 |
| *Bacillus subtilis* PY79 | <0.03[d] |
| *Bacillus mojavensis* RO-H-1 | <0.03[d] |
| *Bacillus spizizenii* DV1-B-1 | <0.03[d] |
| *Bacillus amyloliquefaciens* H | 0.06 |
| *Bacillus clausii* ATCC9799 | <0.03[d] |
| *Pseudomonas aeruginosa* PA14 | <0.03[d] |
| *Serratia marcescens* BS303 | 0.29 |
| *Escherichia coli* MG1655 | 2.10 |
| *Salmonella enterica* serovar Typhimurium LT2 | 1.49 |
| *Salmonella enterica* serovar Typhimurium 14028s | <0.03[d] |

[a] $S7_{50}$ base ($S7_{50}$ salts and metals), to which was added each of the twenty amino acids to a concentration of 3 mM. See Materials and Methods.

[b] Experimental blank

[c] Values from a single culture of each strain are reported.

[d] Concentration below detection limit.

**Table S4.** Strains used in this study.

| Strain | Genotype | Source/Construction[a] |
|---|---|---|
| PY79 | Wild-type | (1) |
| JLG364 | *ΔxylA::*$sspB^{Ec}$ΩCm | (2) |
| JLG366 | *amyE::*P*sspA-*$sspB^{Ec}$ΩCm | (2) |
| JLG3306 | *accA-4x(ala-ser)-ssrA*ΩKm | pJLG468→ PY79 (Km) |
| JLG3329 | *ald-4x(ala-ser)-ssrA*ΩKm | pJLG478→ PY79 (Km) |
| JLG3958 | *yxiA*::*Pc-tomato*ΩKm | pJLG884 → PY79 (Km) |
| JLG3967 | *ald-4x(ala-ser)-ssrA*ΩKm *ΔxylA::*$sspB^{Ec}$ΩCm | JLG3329→ JLG364 (Km) |
| JLG3974 | *yxiA*::*Pc-mNeonGreen*ΩKm | pJLG911 → PY79 (Km) |
| JLG4003 | Δ*ald*::Km | BKK31930 → PY79 (Km) |
| JLG4008 | Δ*ansB*::Km | BKK23570 → PY79 (Km) |
| JLG4014 | Δ*sdaAB*::Km | BKK15850 → PY79 (Km) |
| JLG5146 | Δ*gerAB*::Em | BKE33060 → PY79 (MLS) |
| JLG5148 | Δ*bcd*::Km | BKK24080 → PY79 (Km) |
| JLG5149 | Δ*ald*::Km Δ*gerAB*::Em | JLG5146 → JLG4003 (MLS) |
| JLG5211 | Δ*ansA*::Km | BKK23580 → PY79 (Km) |
| JLG5212 | Δ*hutH*::Km | BKK39350 → PY79 (Km) |
| JLG5213 | Δ*tdh*::Km | BKK16990 → PY79 (Km) |
| JLG5214 | Δ*ald*::Em | BKE31930 → PY79 (MLS) |
| JLG5216 | Δ*ald*::Em *yxiA*::*Pc-tomato*ΩKm | JLG5214 → JLG3958 (MLS) |
| JLG5246 | Δ*gerBB*::Em | BKE35810 → PY79 (MLS) |
| JLG5247 | Δ*gerKB*::Em | BKE03720 → PY79 (MLS) |
| JLG5248 | Δ*ald*::Km Δ*gerBB*::Em | JLG5246 → JLG4003 (MLS) |
| JLG5249 | Δ*ald*::Km Δ*gerKB*::Em | JLG5247 → JLG4003 (MLS) |
| JLG5250 | Δ*bcd*::Km Δ*gerAB*::Em | JLG5148 → JLG5146 (Km) |
| JLG5251 | Δ*bcd*::Km Δ*gerBB*::Em | JLG5148 → JLG5246 (Km) |
| JLG5252 | Δ*bcd*::Km Δ*gerKB*::Em | JLG5148 → JLG5247 (Km) |
| JLG5317 | Δ*bcd*::Em | BKE24080 → PY79 (MLS) |

| | | |
|---|---|---|
| JLG5318 | Δ*bcd*::Em *yxiA*::*Pc-tomato*ΩKm | JLG5317 → JLG3958 (MLS) |
| JLG5337 | *ald-4x(ala-ser)-ssrA*ΩKm *amyE*::P*sspA-sspB*$^{Ec}$ΩCm | JLG3329→ JLG366 (Km) |
| 28A5 | *Bacillus mojavensis* RO-H-1 | *Bacillus* Genetic Stock Center |
| 2A12 | *Bacillus spizizenii* DV1-B-1 | *Bacillus* Genetic Stock Center |
| 3A13 | *Bacillus amyloliquefaciens* H | *Bacillus* Genetic Stock Center |
| 3A15 | *Bacillus clausii* ATCC9799 | *Bacillus* Genetic Stock Center |
| K198 | *Pseudomonas aeruginosa* PA14 | D. Unterweger, MPI for Evol. Biol. |
| K425 | *Serratia marcescens* BS303 | D. Unterweger, MPI for Evol. Biol. |
| CGSC8237 | *Escherichia coli* MG1655 | *Coli* Genetic Stock Center |
| OE60 | *Salmonella enterica* serovar Typhimurium LT2 | D. Unterweger, MPI for Evol. Biol. |
| OE62 | *Salmonella enterica* serovar Typhimurium 14028s | D. Unterweger, MPI for Evol. Biol. |

Plasmid or genomic DNA (left side of the arrow) that was used to transform an existing strain (right side of the arrow) is listed. Antibiotic resistance is indicated parentheses. Km, kanamycin; MLS, erythromycin and lincomycin; Cm, Chloramphenicol.

**Table S5.** Plasmids used in this study.

| Plasmid | Description | Reference |
|---|---|---|
| pJLG3 | pBR329(*ori*)-*ssrA*ΩKm | (3) |
| pJLG24 | pBR329(*ori*)-*Pc-tomato-ssrA*ΩKm | This study |
| pJLG38 | pBR329(*ori*)-*sfGFP*ΩKm | (4) |
| pJLG41 | pBR329(*ori*)-*yxiA*::*Pc-tomato-ssrA*ΩKm | This study |
| pJLG42 | pBR329(*ori*)-*yxiA*::*Pc-tomato*ΩKm | This study |
| pJLG101 | *accA-(ala-ser)-ssrA*ΩKm | (5) |
| pJLG133 | *accA-sfGFP-ssrA*ΩKm | This study |
| pJLG468 | *accA-4x(ala-ser)-ssrA*ΩKm | This study |
| pJLG478 | *ald-4x(ala-ser)-ssrA*ΩKm | This study |
| pJLG884 | pBR329(*ori-bla*)-*yxiA*::*Pc-tomato*ΩKm | This study |
| pJLG911 | pBR329(*ori-bla*)-*yxiA*::*Pc-mNeonGreen*ΩKm | This study |
| pDG1662 | *B. subtilis amyE* integration vector | (6) |

**Detailed descriptions of plasmid construction.**

**pJLG24**. The coding sequence of the *tomato* gene [produced by *in vitro* gene synthesis (Biomatik); sequence provided below] was amplified with primers oJLG36 and oJLG38. The PCR product was restricted with SpeI and SphI and cloned into pJLG3 restricted with NheI and SphI. In the resulting plasmid, *tomato* with the *ssrA* degradation tag at its 3' end is expressed constitutively from the Pc promoter (7). The *tomato-ssrA* gene is linked to a kanamycin resistance gene.

**pJLG41**. Four fragments were joined by Gibson isothermal assembly (New England Biolabs): (i) a fragment of 956 bp containing the first 611 bp of *yxiA* coding sequence and the region immediately upstream of it, amplified from genomic DNA of *B. subtilis* PY79 with primers oJLG65 and oJLG66; (ii) a fragment of 2770 bp encompassing the *Pc-tomato-ssrA* and a kanamycin resistance gene, amplified from pJLG24 with primers oJLG67 and oJLG68; (iii) a fragment of 900 bp including the last 795 bp of the *yxiA* coding sequence and the region immediately downstream of it, amplified from genomic DNA of *B. subtilis* PY79 with primers oJLG63 and oJLG64; (iv) the plasmid backbone containing the pBR329 origin of replication, amplified with primers oJLG69 and oJLG70. The resulting plasmid contains the *Pc-tomato-ssrA*ΩKm construction flanked by sequences with homology to the *yxiA* locus, to enable integration into the chromosome by a double recombination event.

**pJLG42**. pJLG41 was amplified with primers oJLG86 and oJLG87, to generate a PCR product lacking the sequence of the ssrA degradation tag, which was then circularized by ligation. The resulting plasmid contains the *Pc-tomato*ΩKm construct lacking the ssrA degradation tag.

**pJLG133**. sfGFP coding sequence was amplified with primers oJLG55 and oJLG77 from pJLG38 and assembled with pJLG101 amplified with primers oJLG500 and oJLG539 by Gibson assembly.

**pJLG468**. pJLG133 was amplified with primers oJLG1601 and JLG1602 and religated. The primers are phosphorylated at the 5' end, providing substrates for the ligation reaction. After religation, the sfGFP coding sequence is removed, leaving a longer linker encoding four alanines and four serines (ASSASAAS) between the *accA* coding sequence and the ssrA.

**pJLG478**. Four fragments were joined by Gibson isothermal assembly (New England Biolabs): (i) 3' region of the *ald* coding sequence, not including the stop codon, amplified from genomic DNA of *B. subtilis* PY79 with primers oJLG1661 and oJLG1662; (ii) *ssrA*ΩKm fragment amplified from genomic DNA of JLG3306 with primers oJLG7 and oJLG1699; (iii) region immediately downstream of *ald* coding sequence, amplified from genomic DNA of *B. subtilis* PY79 with primers oJLG1663 and oJLG1664; and (iv) a DNA fragment encompassing the spectinomycin resistant gene, the origin of replication, and the beta-lactamase gene from pDG1662, amplified with primers oJLG1696 and oJLG96.

**pJLG884**. Two fragments were joined by Gibson isothermal assembly (New England Biolabs): (i) a *yxiA::Pc-tomato*ΩKm fragment, amplified with primers oJLG1788 and oJLG1789 from pJLG42, and (ii) a fragment containing the origin of replication and ampicillin resistance gene of pBR329, amplified with primers oJLG96 and oJLG1679. The resulting plasmid is similar to pJLG42, but contains an ampicillin resistance gene to facilitate future manipulation.

**pJLG911**. Two fragments were joined by Gibson isothermal assembly (New England Biolabs): (i) a fragment of 735 bp containing the coding sequence of mNeonGreen, generated by gene synthesis (see below, Biomatik) and amplified with primers oJLG1833 and oJLG1834; (ii) an inverse PCR product of pJLG884 that leaves out the *tomato* coding sequence generated by amplification with primers oJLG1827 and oJLG1828. In the resulting plasmid, the *tomato* coding sequence of pJLG884 is replaced by *mNeonGreen*.

*>tomato*
t*aaggagg*atttttagaATGGTGAGCAAAGGCGAAGAAGTTATTAAAGAATTTATGAGATTTA
AAGTTAGAATGGAAGGCTCAATGAATGGCCATGAATTTGAAATTGAAGGCGAAGGCGAAGGC
AGACCGTATGAAGGCACACAAACAGCAAAACTGAAAGTTACAAAAGGCGGCCCGCTGCCGTT
TGCATGGGATATTCTGTCACCGCAATTTATGTATGGCTCAAAAGCATATGTTAAACATCCGG
CAGATATTCCGGATTATAAAAAACTGTCATTTCCGGAAGGCTTTAAATGGGAAAGAGTTATG
AATTTTGAAGATGGCGGCCTGGTTACAGTTACACAAGATTCATCACTGCAAGATGGCACACT
GATTTATAAAGTTAAAATGAGAGGCACAAATTTTCCGCCGGATGGCCCGGTTATGCAAAAAA
AAACAATGGGCTGGGAAGCATCAACAGAAAGACTGTATCCGAGAGATGGCGTTCTGAAAGGC
GAAATTCATCAAGCACTGAAACTGAAAGATGGCGGCCATTATCTGGTGGAATTTAAAACAAT
TTATATGGCAAAAAAACCGGTTCAACTGCCGGGCTATTATTATGTTGATACAAAACTGGATA
TTACATCACATAATGAAGATTATACAATTGTTGAACAATATGAAAGATCAGAAGGCCGACAT
CATCTGTTTCTGTATGGCATGGATGAACTGTATAAATAA

Sequence of the *tomato* gene, produced by gene synthesis. The coding sequence is in upper case. The RBS is in lower case italics.

*>mNeonGreen*
gtgtgt*aaggagg*attttagaATGGTTAGCAAAGGCGAAGAGGATAATATGGCTAGCCTCCC
AGCGACCCACGAACTGCATATTTTTGGCAGCATTAATGGCGTTGACTTTGATATGGTGGGGC
AGGGAACAGGGAACCCTAACGATGGCTATGAGGAGCTCAATCTCAAGAGTACAAAAGGAGAT
TTGCAATTTTCACCTTGGATCCTGGTTCCGCATATTGGCTACGGCTTTCATCAATACTTGCC
TTATCCGGACGGCATGTCCCCGTTCCAAGCTGCGATGGTGGATGGTTCTGGGTACCAGGTGC
ACCGTACTATGCAGTTTGAGGACGGTGCCTCACTGACGGTCAACTATAGATATACTTATGAA
GGCTCACACATTAAGGGTGAGGCCCAAGTTAAAGGAACAGGGTTTCCTGCGGATGGACCGGT
AATGACAAACAGTTTAACCGCTGCGGACTGGTGTCGCTCGAAAAAAACATACCCAAACGATA
AAACGATCATCTCGACCTTCAAATGGAGCTATACTACGGGCAACGGCAAACGCTATCGTTCC
ACAGCACGCACGACTTATACGTTTGCTAAACCGATGGCCGCAAACTACCTCAAAAATCAACC
TATGTACGTGTTCAGAAAAACCGAGTTAAAACATTCAAAAACGGAACTTAATTTTAAAGAGT
GGCAAAAGGCGTTTACAGACGTGATGGGCATGGATGAACTGTATAAATGAtaaatgagagag
gaagaaaac

Sequence of the *mNeonGreen* gene, produced by gene synthesis. The coding sequence is in upper case. The RBS is in lower case italics.

**Table S6.** Primers used in this study.

| Primer | Sequence |
|---|---|
| oJLG7 | AATTGGGACAACTCCAGTG |
| oJLG14 (internal for kanamycin) | ATCGAGCTGTATGCGGAGTG |
| oJLG15 (Internal for kanamycin) | GAAAGAGCCTGATGCACTCC |
| oJLG36 | tttttgcatgcagtt*gttgact*ttatctacaaggtgtgg*cataat*gtgtgTAAGGAGG ATTTTAGAATGGTG |
| oJLG38 | tttttactagtTTTATACAGTTCATCCATGCC |
| oJLG55 | TTTATACAGTTCATCCATGCC |
| oJLG63 | CTTCCTGGACAGGGATATGG |
| oJLG64 | gttaccgtggtaagagccgcGCCTTTAATCTCTCCTCCAG |
| oJLG65 | ctgcaatgataccgcgagacGTGTTGACGGGCAATCAG |
| oJLG66 | AAGCCCGTTTTCGGATTC |
| oJLG67 | atgaatccgaaaacgggcttGCAACGTTGTTGCCATTGC |
| oJLG68 | ccatatccctgtccaggaagTTCCGAATACCGCAAGCG |
| oJLG69 | GCGGCTCTTACCAGCCTAAC |
| oJLG70 | GTCTCGCGGTATCATTGCAG |
| oJLG77 | GCTAGCAGCGCAAGCGC |
| oJLG86 | P-TAAATGAGAGAGGAAGAAAACGG |
| oJLG87 | P-TCATTTATACAGTTCATCCATGCC |
| oJLG96 | GCACTTTTCGGGGAAATGTG |
| oJLG500 | gcgcttgcgctgctagcGTTTACCCCGATATATTGATCTTC |
| oJLG539 | ggcatggatgaactgtataaaGCTAGCGCAGCAAATGATG |
| oJLG1601 | P-GCTAGCGCAGCAAATGATGAAAAC |
| oJLG1602 | P-TGCGCTTGCGCTGCTAGC |
| oJLG1661 | gggttaacgcgtaatccatgCCCGCGGAAAAGTAACAATT |
| oJLG1662 | cttgcgcttgcgctgctagcAGCACCCGCCACAGATGATT |

| | |
|---|---|
| oJLG1663 | cactggagttgtcccaattcTAATTCACAATAAGCTTGCAGA- |
| oJLG1664 | cacatttccccgaaaagtgcCACTTCACGGTTTGCTTCAT |
| oJLG1696 | CATGGATTACGCGTTAACCCAGGTCTAGAGGATCGATCT G |
| oJLG1699 | GCTAGCAGCGCAAGCGCAGCTAGCGCAGCAAATGATG |
| oJLG1679 | catggattacgcgttaacccGCGGCTCTTACCAGCCTAAC |
| oJLG1788 | gggttaacgcgtaatccatgGCCTTTAATCTCTCCTCCAG |
| oJLG1789 | cacatttccccgaaaagtgcGTGTTGACGGGCAATCAG |
| oJLG2106 (*ald* E1) | CGTTGAATTACCGACTCAAGC |
| oJLG2107 (*ald* E2) | GACTTGGAAATGAGGACTGTC |
| oJLG2574 (*bcd* E1) | GATTGCTTTAACAAGCAGAGC |
| oJLG2575 (*bcd* E2) | GGATTAATTGTGAGAATGCG |
| oJLG2576 (*gerAB* E1) | GGATGACTCCATTATCCGTC |
| oJLG2577 (*gerAB* E2) | GATATTCTCACTGTCCCAG |
| oJLG2632 (*hutH* E1) | CAGAAGTTCACAGCATAGC |
| oJLG2633 (*hutH* E2) | CAAGCTCTGTTCCTCTGTTG |
| oJLG2634 (*tdh* E1) | CATATCATGGTACAATCCG |
| oJLG2635 (*tdh* E2) | CATACTATTAAGCTCTGC |
| oJLG2636 (*ansA* E1) | CAACCCATCTTTAGATACAG |
| oJLG2637 (*ansA* E2) | CTTTCTCTTTGCTGGTTATCG |
| oJLG2638 (*sdaAA-AB* E1) | GAAGCAAAGCGCTAATAAG |
| oJLG2639 (*sdaAA-AB* E2) | CATAAATACCGAGTTCATTC |
| oJLG2640 (*ansB* E1) | GATAGCTGATGACGTTGTC |
| oJLG2641 (*ansB* E2) | GTAAACAGCTTATTCATCCG |
| oJLG2644 (*gerKB* E1) | GATCAAACACCAGTTTAAAG |
| oJLG2645 (*gerKB* E2) | CCCTTTTATATAGAAGTGAAC |
| oJLG2646 (*gerBB* E1) | GACACAGTCATCCGTATTC |
| oJLG2647 (*gerBB* E2) | CGATATCTTTGACATCCCAG |

External primers used to verify gene disruptions from the BKK and BKE collections are indicated with the name of the flanked locus followed by E1 or E2.

Primer regions that align to the template are in upper case, homology regions for Gibson assembly in lower case, and Pc promoter -10 and -35 regions in lower case italics.

**Movie S1-S4.** Timelapse phase-contrast microscopy of sporulating wild-type (Movie S1), Ald$^{-}$ (Movie S2), GerA$^{-}$ (Movie S3) and Ald$^{-}$ GerA$^{-}$ (Movie S4) strains. Cells were loaded into a microfluidic chip after growing for 24 hours. Phase-contrast pictures were taken every 5 min for 56 hours. The movies were generated at 30 frames per second. The brightness and contrast of all the movies were adjusted using identical parameters to allow visual comparison.
